# Multi-parameter optical imaging of immune cell activity in chimeric antigen receptor T-cell and checkpoint blockade therapies

**DOI:** 10.1101/2021.10.11.463603

**Authors:** Jinghang Xie, Fadi El Rami, Kaixiang Zhou, Federico Simonetta, Zixin Chen, Xianchuang Zheng, Min Chen, Preethi B. Balakrishnan, Sheng-Yao Dai, Surya Murty, Israt S. Alam, Jeanette Baker, Robert S. Negrin, Sanjiv S. Gambhir, Jianghong Rao

## Abstract

Longitudinal multimodal imaging presents unique opportunities for noninvasive surveillance and prediction of treatment response to cancer immunotherapy. In this work we first designed a novel granzyme B activated self-assembly small molecule, G-SNAT, for quantitative assessment of cytotoxic T lymphocyte mediated cancer cell killing *in vivo*. In lymphoma tumor bearing mice, the retention of cyanine 5 labeled G-SNAT-Cy5 was shown to be highly correlated to CAR T-cell mediated granzyme B release and tumor eradication. In colorectal tumor-bearing transgenic mice, expressing firefly luciferase in hematopoietic cells, and which received combination treatment of anti-PD-1 and anti-CTLA-4, longitudinal bioluminescence and fluorescence imaging revealed the dynamics of immune cell expansion, trafficking, tumor infiltration, and cytotoxic activity which predicted therapeutic outcome before tumor shrinkage was evident. These results support further development of G-SNAT for imaging early immune response to checkpoint blockade and CAR T-cell therapy in patients and highlight the utility of multimodality imaging for improved mechanistic insights into cancer immunotherapy.

## Introduction

Cancer immunotherapy, mainly immune checkpoint blockade and adoptive cell transfer (e.g. Chimeric antigen receptor-CAR T-cell therapy), has significantly improved survival rates in many cancers by providing robust and sustained therapeutic effects (*1, 2*). Yet, many hurdles such as complex immune evasion mechanisms, dysfunction of T lymphocytes, and the immunosuppressive tumor microenvironment (*3–11*) limit treatment efficacy (*12, 13*). Additionally, life threatening immune-related side effects are not uncommon (*14*). Tremendous efforts are now being devoted to understanding and targeting these mechanisms for the development of newer, more effective therapies with improved safety profiles. Non-invasive molecular imaging is a promising strategy for monitoring whole body immune responses both during disease progression and upon treatment induction (*15*).

Radiolabeled antibodies (*16–19*), peptides (*20*), or small molecules (*21, 22*) targeted to surface markers or that show uptake by specific immune cell populations, have been extensively studied for imaging the immune response. Indeed, the presence of specific tumor-infiltrating immune cells for example CD8+ cytotoxic T lymphocytes (CTLs), correlates with a favorable response to checkpoint blockade therapy (*23–25*). Nevertheless, several known immunotolerant mechanisms associated with the suppressive tumor microenvironment such as T-cell anergy, exhaustion or senescence may compromise the accuracy for predicting the therapeutic response based on the mere presence of cell subsets (*26, 27*). In addition to imaging immune cell infiltration, an alternative strategy for monitoring the immune activity responsible for tumor eradication, namely granule-mediated cytotoxicity by natural killer (NK) cells and CTLs has been proposed (*28, 29*). Granule-mediated cytotoxicity, one of the dominant cytotoxic mechanisms against cancerous cells, involves the release of granzyme B (gzmB)-loaded granules to induce lysis of target cells (*30*). Secreted gzmB is intrinsically stable, representing an ideal biological signal of cytotoxicity (*31*). A number of fluorescent protein constructs (*32, 33*) and molecules (*34*) have been proposed using optical imaging to study gzmB function *in vitro*. A handful of fluorogenic (*35–37*), photoacoustic (*37*), chemiluminescent (*29*), and PET (*28*) probes have been developed for imaging gzmB activity or its distribution *in vivo*. These studies focused on probing gzmB in different preclinical models and have supported gzmB as a reliable biomarker of cytotoxic cell activation in anti-cancer responses (*28, 38, 39*).

Given the highly complex and regulated nature of the immune system, multimodal imaging of several aspects of immune response such as immune cell expansion, trafficking, tumor infiltration, and granule-mediated cytotoxicity provides a unique opportunity for longitudinally monitoring the complicated spatiotemporal dynamics of immune activation. In this study, we first developed a sensitive fluorescent imaging probe allowing noninvasive surveillance of granzyme B function *in vivo*. We show that this fluorescent probe, termed a granzyme B sensitive nanoaggregation tracer (G-SNAT), was able to image the activity of granzyme B in models of CAR T-cell and checkpoint blockade therapies. *In vivo* bioluminescence imaging was employed to determine immune cell expansion, trafficking and tumor infiltration. Our results reveal distinct whole-body patterns of immune cell migration and associated cytotoxicity among checkpoint inhibitor treated responder, nonresponder, and non-treated groups. We also discovered that gzmB can maintain partial hydrolytic activity under acidic conditions within cytotoxic granules, and G-SNAT was able to detect gzmB activity within the CTLs. These results highlight the value of multimodal longitudinal imaging for revealing complex biological events and their careful orchestration in response to immunotherapy and further support the development of gzmB imaging for predicting patient response to checkpoint blockade and CAR T-cell therapy.

## Results

### Design of gzmB-sensitive nanoaggregation probes

In effective CAR T-cell or checkpoint blockade therapy (Fig. 1A), CTL extravasate, infiltrate and engage cancer cells to establish immunological synapses which enable rapid, polarized release of cytotoxic granules containing effector proteins perforin and gzmB at the synaptic cleft (Fig. 1B). Perforin, as named, directly perforates target-cell plasma membrane and oligomerizes in a calcium dependent manner into a conduit which allows the passive diffusion of gzmB to trigger programed cell death (Fig. 1C). GzmB is a serine protease that cleaves an IEFD (Ile-Glu-Phe-Asp) peptide motif (preferred substrate in mice) to activate downstream caspase signaling and trigger DNA fragmentation and apoptosis (Fig. 1C) (*40–43*). This unique mechanism in the cytotoxic immune response provides efficient delivery of active gzmB into target cells, in addition to an endocytosis dependent mechanism that has also been proposed (*44, 45*).

**Fig. 1.**
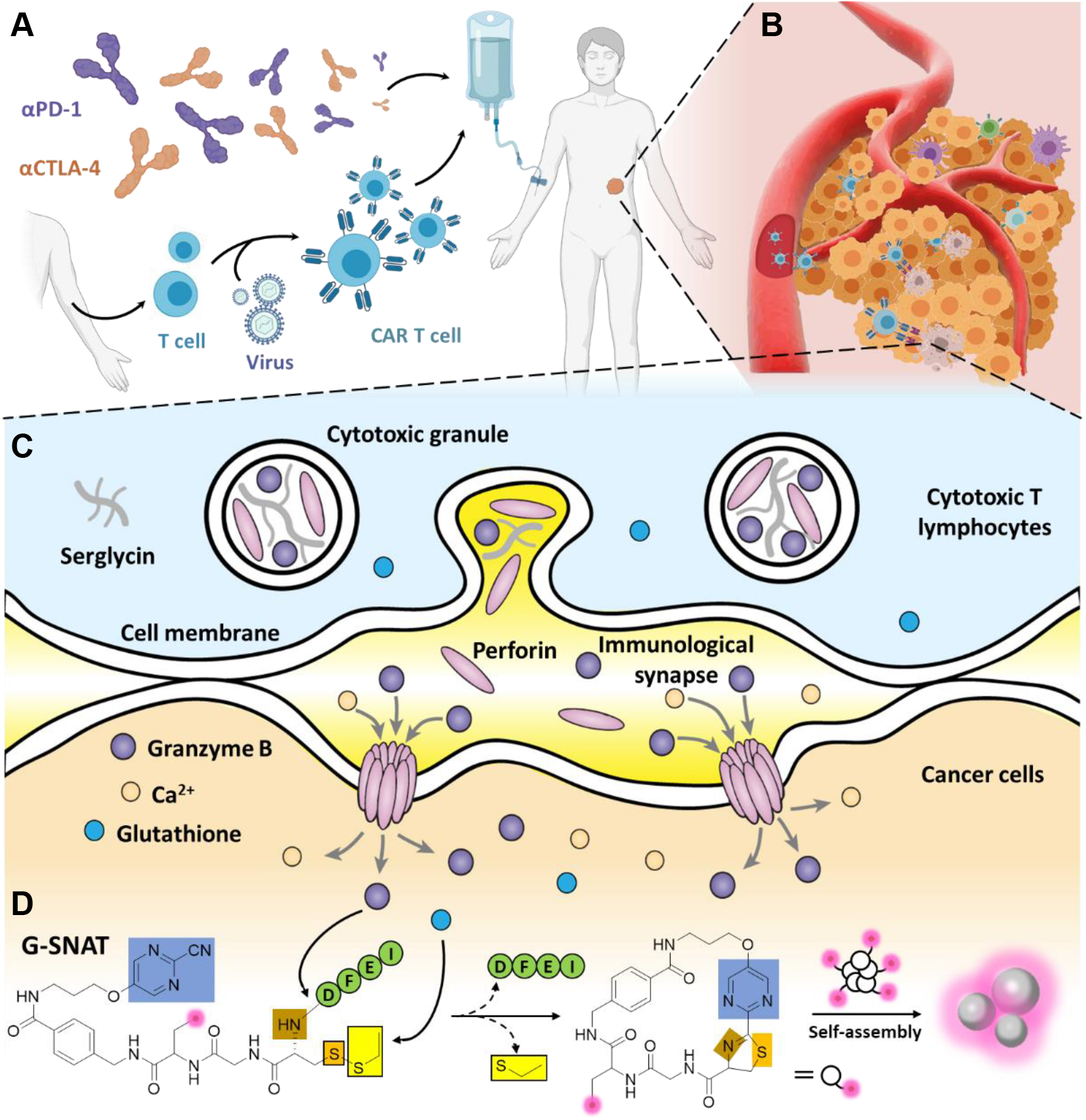
Mechanism of *in vivo* imaging of gzmB activity in tumors by G-SNAT. **A)** Treatment of a patient with checkpoint inhibitors or CAR T-cell therapy. **B)** Effector T cells (blue) can extravasate from blood vessels and infiltrate tumors to kill cancer cells (yellow). **C)** Production of gzmB and perforin by cytotoxic T lymphocytes and their delivery across an immunological synapse into cancer cells. **D)** Proposed gzmB and reduction-controlled conversion of G-SNAT into G-SNAT-cyclized through the biorthogonal intramolecular cyclization, followed by self-assembly into nanoaggregates *in situ*. Blue square, the 2-cyanopyrimidine; brown and orange, amino and thiol groups of D-cysteine, respectively; yellow, thioethyl masking group; green, the capping peptide residues; pink, fluorophore Cy5.

Based on our highly modular TESLA (target enabled *in situ* ligand aggregation) platform (*46*), G-SNAT was designed (Fig. S1 and S2) which uniquely enabled the projection of specific catalytic activity of gzmB during immune response assessment to condensed molecular aggregates. As illustrated in Figure 1D, G-SNAT contains: 1) a 2-cyanopyrimidine group (blue), 2) a cysteine residue coupled to IEFD substrate (green) at the amino group (brown) with a disulfide bond at the mercapto group (orange), and 3) a propargylglycine residue between pyrimidine and the cysteine for labeling with a fluorophore, radioisotope, or other contrast agents (pink). After hydrolysis and cleavage of IEFD by gzmB as well as the reduction of disulfide bond by intracellular glutathione (GSH), the product cyclizes intra-molecularly into macrocyclics which are rigid, hydrophobic, and susceptible to intermolecular interactions that trigger nanoaggregation *in situ* (*46*). For studies *in vitro*, both a cyanine 5 fluorophore (Cy5) pre-conjugated G-SNAT-Cy5 probe (Fig. S2) and post cell culture click reaction at the propargylglycine alkyne handle with an azido cyanine 5 (Cy5, far-red fluorophore) were applied for pinpointing the aggregated G-SNAT in cells.

### Macrocyclization and nanoaggregation of G-SNAT *in vitro*

Incubation of G-SNAT with recombinant mouse gzmB enzyme (1 μg/ml), and tris(2-carboxyethyl)phosphine (TCEP, 2 mM) to mimic the intracellular reducing environment at 37 °C overnight enabled the macrocyclization of G-SNAT (10 μM; retention time, *T*_R_ = 14.2 minutes) to give G-SNAT-cyclized (*T*_R_ = 8.8 minutes), as shown in high-performance liquid chromatography (HPLC) and confirmed by mass spectrometry (Fig. 2A). Since IEFD has been reported to be a preferred substrate of mouse gzmB and its specificity has been validated (*36, 47*), we conducted a kinetic study with caspase 3 as a control, which was believed to share a pool of substrates with gzmB like the nuclear mitotic apparatus protein (NuMA, Val-Leu-Gly-Asp) and DNA-dependent protein kinase catalytic subunit (DNA-PKcs, Asp-Glu-Val-Asp) (*48*). It was shown that gzmB specifically activated G-SNAT (10 μM) and converted it to cyclized product (Fig. 2B and Fig. S3). The assembled nanoaggregation was imaged with a transmission electron microscopy (TEM) and the size distribution was acquired by dynamic light scattering (DLS) (Fig. 2C and 2D). The average diameter of the aggregated nanostructures was 396 nm, ranging from 190 to 955 nm. The nanoaggregates were found nearly neutral by zeta potential analysis (Fig. 2E). Together, these results demonstrate that gzmB specifically hydrolyzes IEFD and induces macrocyclization of the products leading to intermolecular interaction and nano-aggregation. The stability of G-SNAT-Cy5 in mouse serum was evaluated by HPLC, and indicated a half-life of approximately 8-hours (Fig. 2F and Fig. S4).

**Fig 2.**
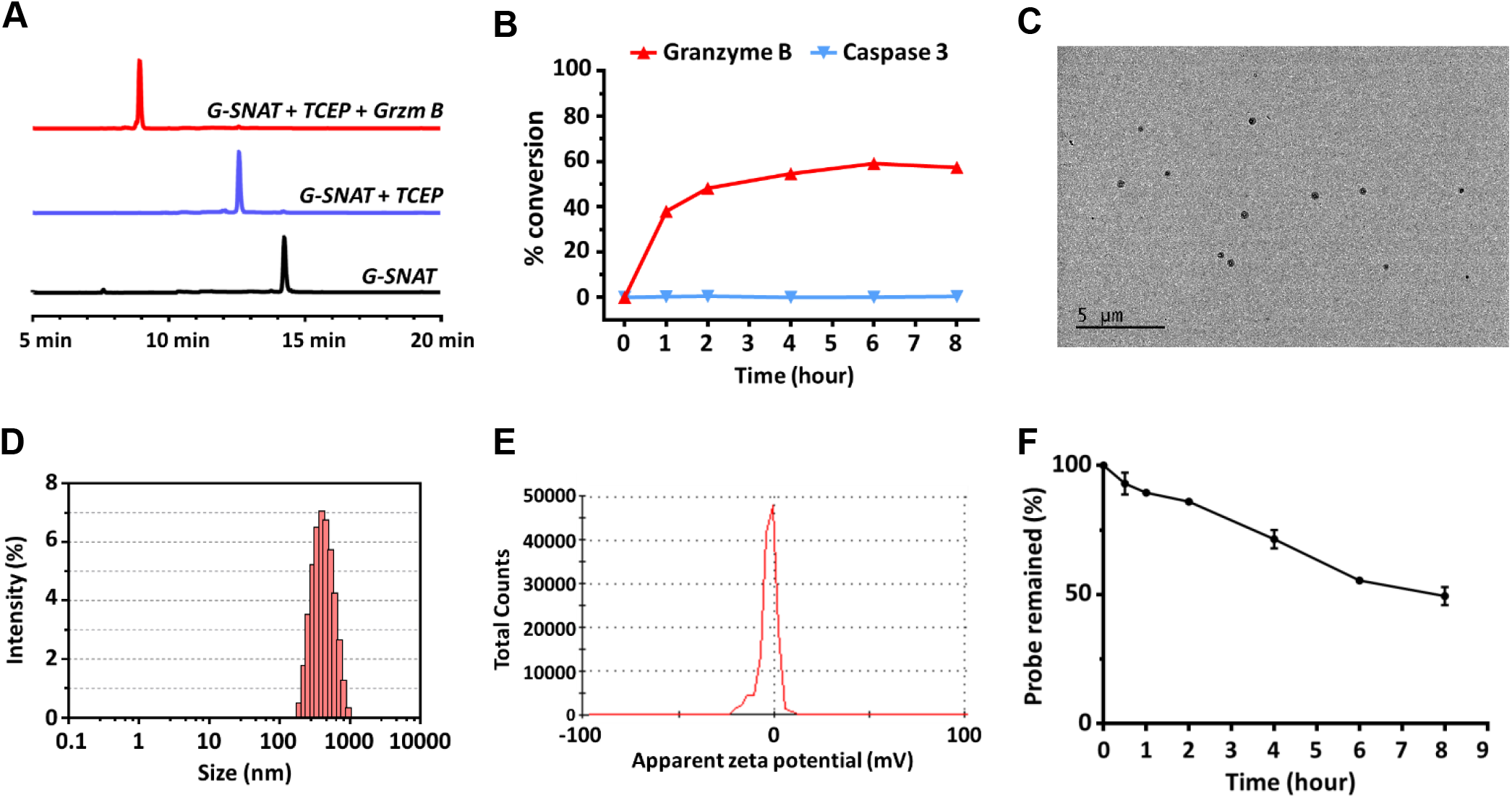
*In vitro* characterization of the G-SNAT probes. **A)** HPLC traces of G-SNAT in gzmB assay buffer (black, T_R_=14.2 minutes) and the incubation of G-SNAT (10 μM) with TCEP (blue, T_R_=12.8 minutes) and recombinant mouse gzmB (1 μg/ml) overnight at 37 °C (red, T_R_=8.9 minutes). **B)** The enzymatic reaction kinetics and specificity studies by longitudinal monitoring of percentage conversion of G-SNAT (10 μM) into G-SNAT-cyclized after incubation with equal amounts (100 U) of recombinant mouse GzmB (0.05 μg/ml) and human caspase-3. **C)** TEM image of nanoaggregates after incubation of G-SNAT (100 μM) with recombinant mouse GzmB (1 μg/ml) overnight at 37 °C in assay buffer. **D)** DLS analysis of diluted (2x) G-SNAT-cyclized (50 μM) showing the distribution of particle sizes. **E)** Measurement of the zeta potential of G-SNAT-cyclized. **F)** Analysis of the stability of G-SNAT in mouse serum by HPLC. The percentage of probe remaining after incubation of 100 μM of G-SNAT-Cy5 in mouse serum at 37 °C for indicated times was obtained by calculating the percentage of peak area (mAU*min) of probe on the corresponding HPLC trace; error bars (S.D.) are calculated from two separated experiments.

### G-SNAT-Cy5 imaging report gzmB activity in cytotoxic T cells and CAR T-cell engaged cancer cells

To examine whether G-SNAT-Cy5 (Fig. 3A) could report gzmB activity in cytotoxic T cells, we isolated CD8+ T cells from BALB/c mice, activated them with anti-CD3/CD28 coated beads, and divided these cells into two groups: one was used to generated CD19-28ζ CAR-T cells using retroviral transduction as previously described (*49*) while the other group was left untransduced. Non-activated naïve T cells (CD44^low^CD62L^high^) were prepared as control (Fig. S5). After 2.5 hours of incubation with G-SNAT-Cy5 (5 μM), confocal microscopic imaging revealed fluorescence in both activated untransduced CD8+ and CAR T cells but not in naïve T cells (Fig. 3B, movies S1, S2 and S3). After its synthesis, gzmB is packaged in lytic granules to prevent self-killing and to facilitate trafficking to the immunological synapse. The images suggested a granular sequestration of intracellular gzmB. To validate that the granular gzmB retained enzymatic activity, we tested recombinant mouse gzmB under acidic (pH5.5) conditions and found that gzmB maintained about 26.6% and 24.3% of its activity at pH5.5 after two- and four-hours incubation with G-SNAT-Cy5 at 37 °C, respectively (Fig. 3C and Fig. S6). Bioluminescence assay with A20 cells expressing firefly luciferase (A20^Luc+^) confirmed that the cytotoxic function of these CAR T cells was not affected by incubation with G-SNAT or G-SNAT-Cy5 overnight (Fig. 3D and Fig. S7). Further cell viability study showed that both G-SNAT and G-SNAT-Cy5 were well tolerated by A20^Luc+^, CD8+ T and CAR T cells (Fig. S8).

**Fig 3.**
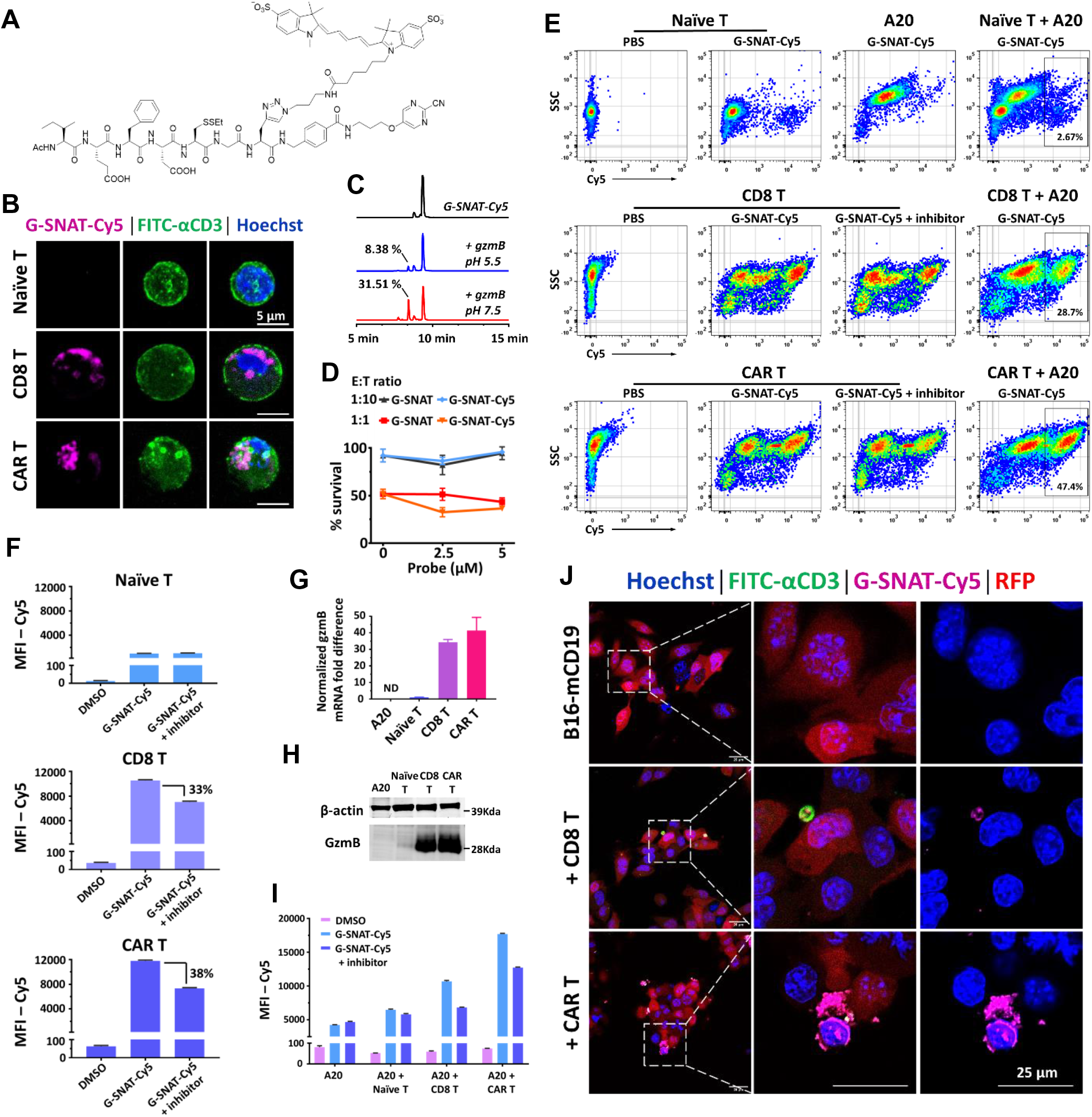
Imaging of gzmB activity in naïve CD8+, activated untransduced CD8+, CD19-28ζ CAR T cells and cancer cells. **A)** Structure of G-SNAT-Cy5. **B)** Microscopic imaging of T cells incubated with G-SNAT-Cy5 (5 μM) for 2.5 h then stained with Hoechst (blue, Ex390/Em440) and FITC conjugated CD3 antibody (green, Ex488/Em520). Magenta (Ex650/Em670) represents retained G-SNAT-Cy5. Scale bar indicates 5 μm. **C)** HPLC traces of G-SNAT-Cy5 (black), and incubation with gzmB (0.05 μg/ml) in MES buffer (pH5.5, blue) or assay buffer (pH7.5, red) at 37 °C for 2 h; 8.38% and 31.51% indicate the percent conversion relative to G-SNAT-Cy5 peak as calculated from the peak areas (mAU*min). **D)** G-SNAT or G-SNAT-Cy5 (0, 2.5 and 5 μM) showed no impact on the cytotoxic function of CD19-28ζ CAR T cells against A20^Luc+^ cells. **E)** Flow cytometry analysis of G-SNAT-Cy5 treated naïve, activated untransduced CD8+ and CAR T cells with or without a GzmB inhibitor or A20 cells. 10,000 cells were analyzed in T cell groups; 20,000 cells were analyzed in T cells incubated with A20 cancer cells at a 1: 1 ratio. The mean fluorescent intensity (MFI) in deep red (Cy5, Ex650, Y axis) were acquired, analyzed, and plotted by Flowjo. **F)** The MFI-Cy5 signals of T cells in **E)** were plotted and compared to show percentage inhibition. Error bars represent standard deviation. **G)** Quantitative RT-PCR analysis of gzmB. ND-nondetectable. **H)** Western blot analysis of gzmB in cell lysate (25 μg). **I)** Flow cytometry analysis of DMSO, G-SNAT-Cy5 (5 μM) or a gzmB inhibitor treated A20 cells incubated with PBS, naïve, activated untransduced CD8+ or CAR T cells in suspension at a 1: 1 ratio for 2.5 h shown in the right panel of **E)** were plotted. Error bars represent standard deviation. **J)** Activated untransduced CD8+ or CAR T cells treated B16-mCD19 cells at a 2: 1 ratio in the presence of G-SNAT-Cy5 probe (5 μM) for 3.5 h. Cells were gently washed, stained with Hoechst (blue, Ex390/Em440) and FITC conjugated CD3 antibody (green, Ex488/Em520), fixed and mounted for confocal microscope to show the stacked image. Magenta (Ex650/Em670) represents retained G-SNAT-Cy5. Red represents RFP (Ex550/Em580). Scale bars indicate 25 μm.

An interesting observation was that gzmB-packed granules displayed a denser fluorescent cluster in CAR T versus untransduced activated CD8+ T cells (Fig. 3B). This might concur with faster lytic granule recruitment to nonclassical CAR T-cell immune synapses described by Davenport (*50*), characterized by lack of Lck clustering at small immunological synapses, which was inherently different than the Lck-rich, large immunological synapses in wild type T cell. To prove that the intracellular activation of G-SNAT-Cy5 is specific to gzmB, we included a competitive gzmB inhibitor (Ac-IETD-CHO) (*51*) and quantified the Cy5 fluorescent signal with flow cytometry (Fig. 3E and F). On average, CAR T cells presented more than 10-fold elevation in Cy5 fluorescent intensity over naïve T cells, although a small portion of selected CD8+ T cells remained in naïve T cells after pan selection. For activated untransduced CD8+ T cells, it was elevated by around 9-fold (Fig. 3F). When gzmB inhibitor was added, the mean fluorescence intensity (MFI-Cy5) in CAR T cells dropped by 38% and 33% in activated untransduced CD8+ T cells (Fig. 3F). Collectively, these results demonstrate specific activation and retention of G-SNAT-Cy5 by gzmB in the granules of cytotoxic T cells in a bioorthogonal manner.

We further investigated whether G-SNAT-Cy5 could report the cytotoxic activity in three established CAR T-cell therapy models *in vitro*: CD19-28ζ CAR T cells co-cultured with A20 murine B cell lymphoma cells naturally expressing CD19 and B16 melanoma cells engineered to stably express RFP (red fluorescent protein) and murine CD19 antigen (B16-RFP/mCD19); GD2-4-1BBζ CAR T cells transduced from total T cells co-cultured with SB28 murine glioblastoma cells engineered to stably express RFP and mouse GD2 (SB28-RFP/GD2). Both CAR T cells were freshly prepared and maintained according to previously reported protocols (*49, 52*). Activated untransduced CD8+ T cells and CD19-28ζ CAR-T cells were first incubated with A20 cells in suspension at a 1:1 ratio (Effector: Target) for 2.5 hours. We analyzed naïve, activated untransduced CD8+ and CAR T cells with or without A20 by flow cytometry (Fig. 3E) and quantified the MFI-Cy5. As displayed in two dimentional plots (side scatter-y-axis; Cy5-x-axis) (Fig. 3E), compared to naïve (2.67%) or untransduced CD8+ T cells (28.7%), a dramatic cell population shift along the x-axis was observed in CAR T cells exposed to A20 cells (47.4%), which suggested a higher activation and retention of G-SNAT-Cy5 by gzmB mediated cytotoxic killing upon exposure to CD19 antigen. We analyzed the expression of gzmB in these cancer and T cells at both transcriptional (Fig. 3G) and protein levels (Fig. 3H). GzmB protein was robustly expressed in activated CD8+ and CAR T cells, negligible in naïve T cells, and completely non-detectable in A20 cancer cells. CD19-28ζ CAR transduced T cells expressed more gzmB than the activated untransduced CD8+ T cells, which correlated to the higher retention of G-SNAT-Cy5. When gzmB inhibitor was co-incubated, the overall probe retention also decreased in both activated untransduced CD8+ and CAR T cells treated cancer cells (Fig. 3I). To better image the gzmB mediated cytotoxic cancer killing, we tested adherent B16-RFP/mCD19 cells. Compared to A20 cells that were known for a naturally high expression of CD19, B16-RFP/mCD19 was genetically engineered. We thus adjusted the E:T ratio to 2:1 and incubated for 3.5 hours in the presence of 5 μM of G-SNAT-Cy5. As expected, cyclized and aggregated products were only observed in CAR T cell-engaged cancer cells presenting morphological hallmarks of apoptosis: shrinkage, fragmentation, and package of cell content (Fig. 3J and Fig. S9). Consistent with above T cell imaging results, probe retention was also found in both activated untransduced CD8+ and CAR T cells.

To test the effect of Cy5 on the cell uptake of G-SNAT-Cy5, we repeated the study in GD2-4-1BBζ CAR T-cell treated SB28-RFP/GD2 cells with a post-click imaging strategy as previously described to visualize nanoaggregates after cell incubation (Fig. S10) (*53*). A caspase-3 sensitive nanoaggregation probe C-SNAT4 developed previously was included to detect apoptosis triggered by gzmB. Consistent with pre-labeled probes, G-SNAT was highly activated and retained in CAR T and engaged cancer cells after 2.5 hours incubation at a 2:1 ratio. Additional caspase-3 imaging pinpointed the apoptotic cells without nonspecific detection of gzmB (Fig. S11). Together, these results demonstrate that G-SNAT probe can image the activity of gzmB from CTLs during immunotherapy.

### Bioluminescence and G-SNAT-Cy5 fluorescence imaging reveal gzmB mediated cytotoxic activity in CAR T-cell treated lymphoma models

To assess the ability of G-SNAT-Cy5, to report gzmB activity during CAR T-cell therapy *in vivo*, we first tested a gzmB releasing model by subcutaneously (s.c.) implanting A20 lymphoma cells into the upper right flank of BALB/c Rag2^−/−^γc^−/−^ mice and treated them with CD19-28ζ CAR T ^Luc+^ cells (Fig. 4A). These CAR-T^Luc+^ cells were generated by retroviral transduction of activated CD8+ T cells from transgenic Fluc+ BALB/c mice with hematopoietic cells engineered to stably express firefly luciferase (*54*). When tumor volume reached ∼300 mm^3^, 6 × 10^6^ CAR T^Luc+^ cells were intratumorally injected. The viability and distribution of these T cells were monitored by bioluminescence imaging with standard intraperitoneal (i.p.) administration of D-luciferin (3 mg) at 24 (day 12) and 48 hours (day 13) post T cell injection. Associated gzmB activity was longitudinally imaged at 1, 2, 4 and 20 hours post tail vein administration of G-SNAT-Cy5 (5 nmol) at day 12 (Fig. 4A). At day 13, the tumors were removed, imaged, and analyzed by western blot and immunofluorescent staining after another round of G-SNAT-Cy5 imaging. As shown in Figure 4B, CAR T^Luc+^ cells maintained good viability post intratumoral injection, although a decay was noticed at day 13 (Fig. S12). The region of interest (ROI) on tumors was defined to quantify the fluorescence intensity of G-SNAT-Cy5 (Fig. S13). The highest MFI-Cy5 was observed at 1 hour post injection in CAR T^Luc+^ cells treated tumors which was averagely 3.1-fold of the PBS treated tumors (Fig. 4C). In *ex vivo* analysis, the detection of G-SNAT-Cy5 highly correlated to the expression of gzmB in tumors (Fig. 4D and E), and the colocalization of CD8α, gzmB and G-SNAT-Cy5 was confirmed by immunofluorescent staining (Fig. 4F).

**Fig 4.**
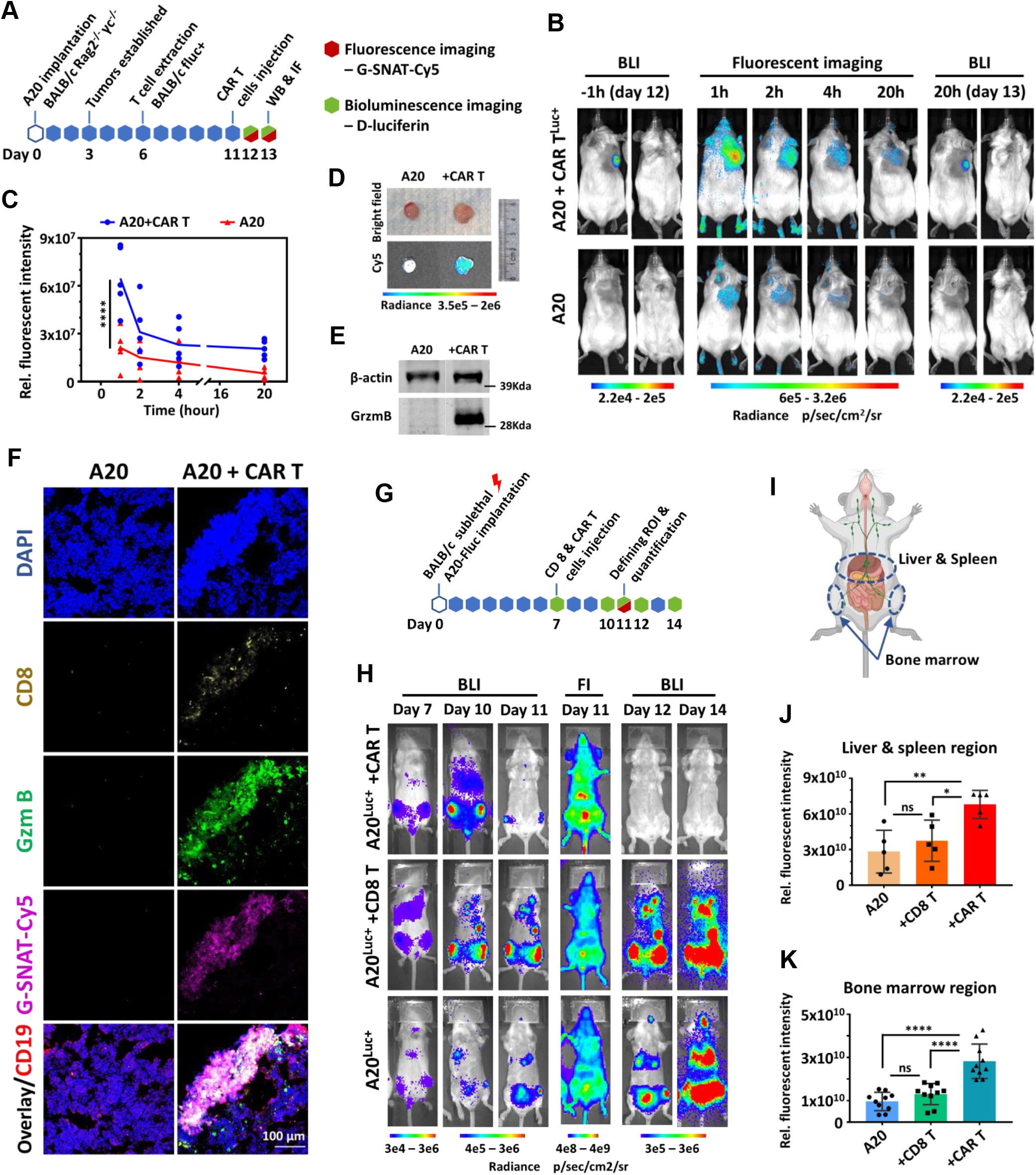
Multimodal optical imaging with G-SNAT-Cy5 and D-luciferin predict lympoid tumor reponse to CAR T-cell therapy. **A)** Illustration of the workflow to generate the subcutaneous A20 lymphoid tumor/CD19-28ζ CAR T^Luc+^-cell therapy model and imaging study with D-luciferin and G-SNAT-Cy5. **B)** Longitudinal bright field, bioluminescence, and fluorescence imaging with D-luciferin and G-SNAT-Cy5 (5 nmol, Ex640/Em690) of A20 implanted (bottom) and CAR T ^Luc+^ cells (top) treated tumor-bearing mice. Representative mice were shown here while full panels were shown in Fig. S12 and S13. **C)** A comparison of the relative fluorescence intensity acquired by defining the ROI on A20 and CAR T ^Luc+^ cells treated tumors, \*\*\*\**p* < 0.0001. **D)** Bright filed and fluorescence imaging of A20 and CAR T ^Luc+^ cells treated tumors at day 13. Ruler unit is cm. **E)** Western blot analysis of gzmB in tumor lysate (50 μg). **F)** Immunofluorescent staining analysis of the tumors in **D)**. **G)** Illustration of the workflow to generate the systemic lymphoma model and imaging study with D-luciferin and G-SNAT-Cy5. **H)** Longitudinal bright field, bioluminescence, and fluorescence imaging with D-luciferin and G-SNAT-Cy5 (5 nmol, Ex650/Em670) of A20^Fluc+^ implanted (bottom), activated untransduced CD8+ T cells (middle) or CAR T cells (top) treated tumor-bearing mice. Representative mice were shown here while full panels were shown in Fig. S14. **I)** Cartoon illustrating the ROI on liver and spleen (middle circle) and bone marrow (lower two circles) regions. **J)** A comparison of the relative fluorescence intensity acquired by defining the ROI on liver and spleen regions of A20, activated untransduced CD8+ or CAR T ^Luc+^ cells treated tumor-bearing mice. \**p* < 0.0332; \*\**p* < 0.0021, ns: not significant. **K)** A comparison of the relative fluorescence intensity acquired by defining the ROI on bone marrow regions of A20, activated untransduced CD8+ or CAR T ^Luc+^ cells treated tumor-bearing mice. \*\*\*\**p* < 0.0001.

To better mimic the clinical scenario, we employed a systemic lymphoma model in which A20^Luc+^ cells were intravenously injected (i.v., tail vein) into sublethally (4.4 Gy) irradiated BALB/c mice (Fig. 4G). In agreement with what has been reported previously (*55*), by the time of CD19-28ζ CAR T-cell administration (2.4 × 10^6^, i.v., retro orbital), 7 days after tumor inoculation, A20^Luc+^ cells were infiltrating the liver, the lymphoid organs, including bone marrow (Fig. 4H, I and Fig. S14). Longitudinal bioluminescence imaging with D-luciferin revealed the therapeutic effect of CAR T cells. Fluorescence imaging performed one hour post tail vein administration (i.v.) of 5 nmol G-SNAT-Cy5 at day 11 showed a significantly higher retention of the probe in both liver, spleen and the bone marrow of CAR T cell treated mice (Fig. 4J and K). Collectively, these results suggest that G-SNAT-Cy5 might be useful in reporting the degree of cytotoxicity in tumors responded to CAR T cell-therapy.

### Bioluminescence and G-SNAT-Cy5 fluorescence imaging monitor tumor response to checkpoint blockade therapy

To visualize the immune activation to checkpoint blockade therapy, we performed longitudinal bioluminescence and fluorescence imaging of a syngeneic mouse model treated with a combined regimen of anti-PD1 and anti-CTLA4 antibodies. As outlined in Figure 5A, transgenic Fluc+ BALB/c mice expressing firefly luciferase in hematopoietic cells were implanted subcutaneously with 1×10^6^ CT26 murine colorectal cancer cells and treated with 200 μg of anti-PD1 and 100 μg of anti-CTLA4 intraperitoneally at day 9, 12, and 15 post inoculation. As indicated by tumor growth curves, this combination therapy triggered a heterogeneous response (Fig. 5B). We defined those treated without complete tumor regression at endpoint (day 35) as nonresponders (grey) and imaged all cohorts (treated responder, nonresponder, and nontreated) longitudinally with D-luciferin, G-SNAT-Cy5 and a previously developed gzmB specific bioluminogenic substrate GBLI2 (Fig. S15) at day 9, 12, 15, 18, and 20. With D-luciferin, the expansion and whole-body trafficking of hematopoietic cells in response to checkpoint inhibitors could be monitored. Additional GBLI2 imaging provided a reference of gzmB activity for validation of G-SNAT-Cy5.

**Fig 5.**
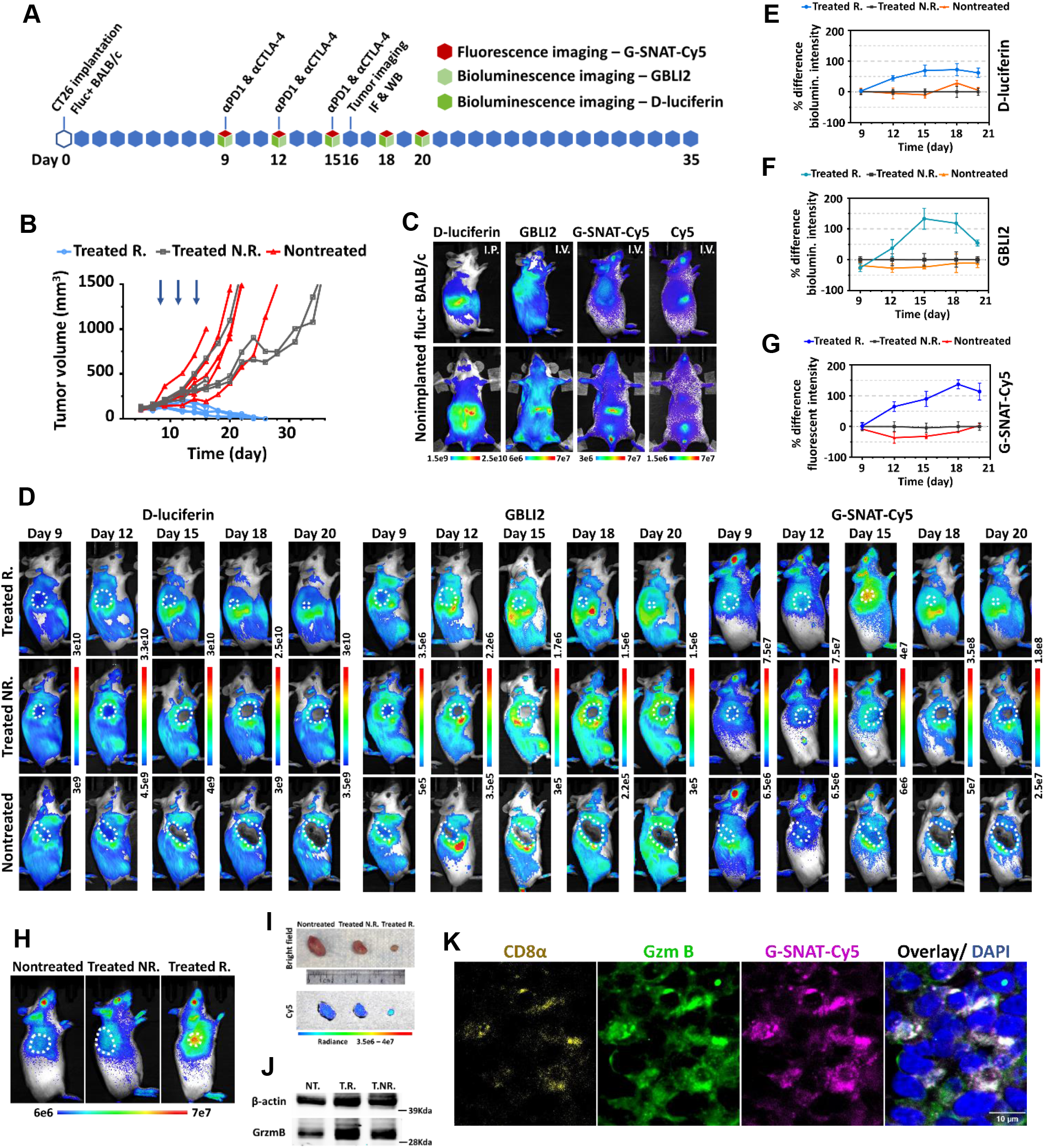
Multimodal optical imaging with G-SNAT-Cy5, GBLI2 and D-luciferin predict colorectal tumor reponse to checkpoint blockade therapy. **A)** Illustration of the workflow to generate the subcutaneous CT26 colorectal tumor in Fluc+ BALB/c mice treated with anti-PD-1 and anti-CTLA-4 and imaging studies with G-SNAT-Cy5, GBLI2 and D-luciferin. **B)** Growth of the tumors in treated responder (Treated R.), treated non-responder (Treated N.R.), and nontreated cohorts. Some of the mice were sacrificed at day 16 for *ex vivo* analysis. **C)** Bright field, bioluminescence, and fluorescence imaging of nonimplanted healthy Fluc+ BALB/c mice showing the base line uptake and distribution of D-luciferin (3 mg, i.p), GBLI2 (200 μg, i.v. retro orbital), G-SNAT-Cy5 (5 nmol, i.v. retro orbital, Ex640/Em690) and Cy5 (5 nmol, i.v. retro orbital, Ex640/Em690). **D)** Longitudinal bioluminescence imaging with D-luciferin (3 mg, i.p.), GBLI2 (200 μg, i.v. retro orbital), and fluorescence imaging with G-SNAT-Cy5 (5 nmol, i.v. retro orbital) of nontreated (bottom), treated non-responder (middle) and responder (top) groups. White circles indicate tumors. Representative mice from each group were shown here. **E)**, **F)** and **G)** Relative bioluminescent and fluorescent intensity of tumors at day 9, 12, 15, 18, and 20 were quantified by defining the ROI on tumors. The percent difference of bioluminescence and fluorescence intensity among treated responder, treated non-responder, and nontreated over the course of imaging were plotted using treated non-responder as the baseline; n=5 for treated R., n=4 for Treated N.R., and n=5 for Nontreated. **H)** Fluorescence imaging with G-SNAT-Cy5 of the mice at day 16 before euthanasia and tumor collection. White circles indicate tumors. **I)** Bright field and fluorescence imaging of tumors collected from **H).** Ruler unit is cm. **J)** Western blot analysis of gzmB. 50 μg tumor lysate was load for analysis. **K**) Immunofluorescence staining analysis of treated responded tumor from **I).** Scale bar indicates 10 μm.

The base line uptake and biodistribution of these probes were first evaluated with non-tumor bearing healthy Fluc+ BALB/c mice (Fig. 5C). Via i.p. injection (3 mg), the bioluminescence from D-luciferin was mainly restricted to the spleen and lower abdomen, which is ideal for imaging immune cell trafficking to the tumor site. GBLI2 imaging (200 μg, i.v. retro-orbital) gave approximately two magnitude weaker bioluminescence but highly concentrated in the spleen (Fig. 5C), to a less extent in the bone marrow, which suggested the major homing organs of CTLs. G-SNAT-Cy5, one hour post i.v. injection (5 nmol, retro orbital), showed a similar preferential activation and retention in the spleen, in contrast to the fast clearance of Cy5 fluorophore alone (5 nmol, i.v. retro orbital) (Fig. 5C). Next, we imaged tumor-implanted mice receiving checkpoint blockade therapy. Consistent with growth curves, bright field imaging showed divergent tumor responses to checkpoint blockade (Fig. 5D and Fig. S16). Responding tumors initially showed an increase in volume at day 12, 3 days post the first treatment, then a sustained shrinkage from day 15 to 20. For treated nonresponding and untreated tumors, continuous growth was observed. When all these cohorts were imaged with D-luciferin, an initial activation and expansion of immune cells were seen in treated cohorts at day 12, indicated by enhanced bioluminescence in the spleen and surrounding tumor. GBLI2 imaging showed the activation of CTLs in all three cohorts regardless of treatment but only the treated responders had obvious CTLs infiltration. By longitudinal imaging, a distinct pattern of gradual immune cell exclusion from tumors appeared in nonresponding and nontreated cohorts. At day 18 and 20, a later stage of tumor eradication in responders, we noticed a large portion of CTLs were homing back to the spleen, whereas the nonresponded and nontreated cohorts still had most of the immune cells surrounding tumors. In comparison to D-luciferin and GBLI2, G-SNAT-Cy5 imaging revealed overall similar distribution of CTLs in responding to therapy at both early (day 9-12) and late stage (day 18-20), but showed greatly elevated and confined fluorescence signal within the tumor at day 15 (Fig. 5D). This result indicated robust gzmB releasing and cytotoxic killing following immune cells tumor infiltration in responders and featured our design of self-nanoaggregation in the detection of gzmB activity *in vivo*.

To quantify the dynamic change of bioluminescent and fluorescent signals, we defined ROIs on tumors and calculated the relative percent difference using treated nonresponded tumors as the reference (Fig. 5E, F, G and Fig. S17). By tracking total immune cells with D-luciferin, an elevated tumor infiltration was observed since day 12, and the infiltrates were stably maintained through the whole process of tumor eradication in responders. GBLI2 and G-SNAT-Cy5 correlated well to immune cell infiltration in responded tumors since day 12 which predicted the therapeutic outcome prior to tumor volume divergence, excepted the continuously elevated signals peaked at day 15 for GBLI2 and day 18 for G-SNAT-Cy5. This data suggested a sustained expression and releasing of gzmB in a relatively stable number of immune cells including CTLs within the tumor. Follow-up mouse and tumor imaging at day 16 confirmed a higher G-SNAT-Cy5 retention in responded tumors (Fig. 5H and I) that correlated well to the level of gzmB expression (Fig. 5J) and localization (Fig. 5K and Fig. S18). Clearly, elevated gzmB afforded better retention of G-SNAT-Cy5 which supported the mechanism of gzmB dependent G-SNAT activation and aggregation. Taken together, by visualizing the dynamics of immune cell expansion, trafficking and gzmB mediated cytotoxicity, we observed the essential role of early and sustained CTLs infiltration in battling solid tumor. G-SNAT-Cy5 gzmB imaging could very well differentiate and predict tumor response to checkpoint blockade therapy.

## Discussion

Despite the influential contributions of immunotherapy for the treatment of diverse cancer types, major roadblocks to basic mechanistic understanding and clinical management are yet to be tackled. Conventional immune profiling relying on tissue, single cell, and molecular level analysis of biopsies or samples at autopsy has many known limitations (*15*). In addition to anatomical imaging, molecular imaging has yielded new strategies to monitor immune activities noninvasively, yet most of the studies have focused on labeling or detecting specific immune cell populations, cytokines, ligands, or other proteins, not adequate in synchronizing dynamics and functions. In this work, we explored multimodal optical imaging for noninvasive and longitudinal measurement of multiple immune parameters in checkpoint blockade and adoptive cell transfer therapy. We developed a novel, gzmB-activated self-assembly small molecule imaging probe G-SNAT, together with luciferase reporter, for tracking immune cell expansion, migration, tumor infiltration, and CTLs mediated cytotoxic cancer killing in Fluc+ transgenic mice.

In the past decade, immunoPET strategy, combining the specificity of monoclonal antibodies and the inherent sensitivity of PET, has dominated immunoimaging and achieved considerable success, such as the translation of ^89^Zr-pertuzumab and ^89^Zr-atezolizumab. As companion diagnostics, immunoPET approaches could assist in tailoring the immunotherapy plan by providing pivotal information regarding the expression and distribution of target antigens prior to and during therapy. Nevertheless, several gaps remain: 1) Membrane associated immune markers are often expressed by more than one cell populations. For instance, CD8 has been mainly identified on CTLs but could also be found on regulatory T cells, natural killer (NK) cells and dendritic cells (*56, 57*), all of which are present in the tumor microenvironment at an early stage (*58*). 2) Expression of immune targets may not fully correlate to therapeutic response such as PD-L1 (*59*). 3) Antibody based imaging probes usually have poor tissue/tumor penetration and a significantly long circulation half-life (days to weeks), which necessitates long half-life radioisotopes associated with high radiation dose (*60*). 4) It is crucial to develop biologically inert antibodies without detrimental immune depletion effects and toxicity (*60*). In comparison, our newly designed G-SNAT may uniquely fill the gaps by targeting specifically the gzmB mediated cytotoxicity. Based on the TESLA strategy which is versatile for multiple imaging modalities and has driven the successful development of a few small molecule imaging probes (*53, 61–63*) including a caspase-3 sensitive PET tracer now in clinical trial, G-SNAT is initially administered in a format of small molecule, upon conversion *in situ* to nanoparticles, and afford a bioorthogonal sensing of CTLs’ killing behavior.

GzmB is produced in the cytosol as a zymogen with a N-terminal Gly-Glu dipeptide that inhibits the assembly of a functional catalytic triad (*64*). In the Golgi, pro-gzmB is tagged with a mannose 6-phosphate for targeting the cytotoxic granules (*65*). Once packaged into granules, dipeptidyl peptidase I (DPPI, cathepsin C) cleaves the Gly-Glu dipeptide and activates gzmB (*66*). The active enzyme is then deposited on a scaffold of serglycin (*67, 68*) under acidic condition (∼pH5.5) (*69, 70*). It was believed that these mechanisms tightly regulated gzmB function within effector lymphocytes, and gzmB maintained little activity prior to being released from granules, supported by cellular imaging of both CTLs and NK cells with gzmB activatable fluorescent protein or peptide-fluorophore constructs which reports that packed gzmB is not active (*32, 33*). However, there are also reports that gzmB could remain partially active in granules and turn on fluorogenic small (*34*) or cell permeable macromolecules (*36*). To confirm this activity under acidic condition, we tested gzmB processing of G-SNAT-Cy5 in MES buffer at ∼pH5.5. This buffer system mimicked the pH condition in lytic granules and has been used for DPPI to process and activate pro-gzmB. We confirmed that gzmB could maintain ∼25% of its activity under the acidic condition, consistent with an early study (*71*). Taking into account that granule-released, serglycin-bound gzmB could trigger apoptosis of target cells as efficient as free enzymes (*72*), we believe the activation and retention of G-SNAT-Cy5 within CTLs are dependent on the gzmB activity. The different results may be explained by the varied permeability of probes to cell or granule membranes that led to distinct pattern of gzmB activity.

To date, five CAR T-cell therapies have been approved by FDA to treat several types of lymphomas and leukemias, as well as multiple myeloma. Despite great success in hematological malignancies, harnessing CAR T cells for treating solid tumors has seen little progress. Current research focuses on improving tumor infiltration, persistence, and potency in heterogeneous and immunosuppressive tumor microenvironment (*73*). As demonstrated in our preclinical lymphoma tumor model, G-SNAT gzmB imaging may facilitate these efforts by reporting the distribution of CAR T cells and the onset of cytotoxic killing. Moreover, this sensitive imaging technology may assist in the detection of “on-target, off-tumor” effects and allow earlier intervention, to mitigate immune-related toxicity. In the subcutaneously implanted, semi-solid lymphoid tumors, a decrease of CAR T cell viability post intratumoral injection was noticed with bioluminescence imaging (Fig. 4B and Fig. S12). This decay might reflect the gradual exhaustion of CAR T cells overwhelmed by CD19 antigens within the tumor microenvironment. On the other hand, A20 lymphoma cells were known to express high level of PD-L1 and a combined treatment of anti-PD-L1 antibody and ibrutinib, an approved Bruton’s tyrosine kinase inhibitor, could cure established A20 tumors (*74*). It is worth future investigation how these CAR T cells lost persistence within the semi-solid A20 tumors and whether checkpoint inhibitor could reverse that. It will also be interesting to examine if G-SNAT gzmB imaging could be utilized to differentiate dysfunction CAR T cells *in vivo* since cumulative data have suggested that senescent T cells have reduced expression of gzmB (*75*) and exhausted CTLs have impaired granzyme packaging and degranulation (*76, 77*) thus defect in gzmB releasing and cytotoxicity.

While checkpoint inhibitors have been adopted to treat a variety of solid tumor types, only a small percentage of patients exhibit complete response, and early detection of anti-tumor immunity remains challenging. By imaging CT26 colorectal tumors whose response to anti-PD-1/anti-CTLA4 were dichotomous, we found that checkpoint blockade therapy cured cancer by stimulating and maintaining early immune cell infiltration including activated and expanded CTLs, which led to gradually intensified cytotoxicity at tumor site that was inversely proportional to tumor volume (Fig. 5). These findings are consistent with a previous immune profiling by flow cytometry and gene expression analysis showing responded CT26 tumors contained expanded T and NK cell populations with CD8+ T cells nearly doubled three days after the third round of anti-PD-1/anti-CTLA-4 treatment (*78*). At this point, G-SNAT or GBLI2 imaging is not able to differentiate gzmB activity from NK and certain gzmB expressing regulatory T cells or between newly infiltrated and tumor surrounding effector lymphocytes, but unlike conventional immune profiling, G-SNAT gzmB imaging would advance spatially and temporally correlated surveillance of cytotoxicity and predict tumor early response *in vivo*. Through longitudinal imaging, we observed a pattern of so called “immune dessert” in both advanced nontreated and treated nonresponded tumors. With current imaging, we could not rule out the possibility that infiltrated immune cells or CTLs slowly lost their viability *in situ*. But a similar pattern displayed by longitudinal immunoPET with an ^89^Zr-labeled PEGylated variable region segment of camelid heavy chain-only antibody targeting CD8 from Rashidian, et al (*18*) supported our conclusion that CTLs were exiled from advanced tumors. Future hybridization of G-SNAT functional imaging and cell tracking immunoPET should sketch a more comprehensive picture of how immune system interacts with tumor cells upon checkpoint blockade and may generate new insights leading to the development of more potent therapies.

As bioluminescence imaging of total immune cells requires pre-introduction of luciferase and immunoPET targeting general immune markers such as CD45 (*79*) has shown high overall signal through the body, it seems not feasible to predict immunotherapy in patients by tracking total immune cells. Multimodal MRI imaging with hyperpolarized ^13^C pyruvate and fumarate for detecting tumor metabolic and physiologic changes during checkpoint blockade has seen progress yet to be optimized for translation (*80*). While gzmB inhibitor based ^68^Ga-NOTA-GZP and ^18^F-AIF-mNOTA-GZP probes have shown great value for assessing immunotherapy response (*28, 38, 39, 81*), G-SNAT-Cy5 might provide a convenient alternative to monitor and predict tumor response with optical imaging in preclinical animal models. Future labeling of G-SNAT with a PET radioisotope will allow direct comparison of different gzmB imaging strategies.

In conclusion, we developed a novel gzmB activated self-assembly imaging probe and proved conceptually the G-SNAT gzmB imaging could be utilized to monitor cytotoxic activity and predict tumor response to CAR T-cell and checkpoint blockade therapies. Together with longitudinally whole-body bioluminescence imaging, the activation, expansion, trafficking and tumor infiltration of total splenocytes or CTLs in responding to immunotherapies were visualized, which revealed an essential role of early immune infiltration and persistent cytotoxic activity in curing cancer. These results support further development of G-SNAT for PET imaging of immune response in patients and offer new opportunities for studying the cytotoxic function in the context of immune cell interplay, tumor microenvironment, and new cancer immunotherapy.

## Acknowledgments

We thank Stanford Neuroscience Microscopy Service (NMS) (supported by NIH NS069375) for help on confocal imaging, Stanford Shared FACS Facility for instrumentation and assistance with flow cytometry, Functional Genomics Facility at Stanford for access to LI-COR Odyssey for Western blots, and Stanford Center for Innovation in In-Vivo Imaging (SCI^3^) for instrumentation and assistance with animal fluorescent imaging.

## Author contributions

J.X., F.E.R and J.R. designed and led the study. K.Z., Z.C. S-Y.D. and J.X. performed the probe synthesis and analysis. X.Z. performed TEM and DLS studies. F.E.R. and S.M. generated activated untransduced CD8+ and CAR T cells. J.X. performed confocal imaging. F.S., F.E.R. and J.X. generated and imaged the CAR T-cell therapy mouse models. M.C., P.B.B. and J.X. generated and imaged the checkpoint blockade therapy mouse model. P.B.B. performed immunofluorescence staining. F.S., I.S.A. J.B., R.S.N., and S.S.G. contributed to the design and data analysis of CAR T-cell therapy models. J.B., F.S. and R.S.N. provided transgenic and Rag2^−/−^γc^−/−^ mice for the study. J.X., F.E.R. and J.R. analyzed all the data, and wrote the manuscript with inputs from all co-authors.

## Data and materials availability

All data associated with this study are present in the paper or Supplementary Materials. Materials are available and will be provided under the material transfer policies of Stanford University. These requests should be directed to the corresponding author.

## SUPPLEMENTARY MATERIALS

Materials and Methods

Fig. S1. Synthesis of G-SNAT precursor.

Fig. S2. Synthesis of G-SNAT and G-SNAT-Cy5.

Fig. S3. Representative HPLC traces in gzmB and caspase-3 kinetic study.

Fig. S4. Representative HPLC traces of G-SNAT-Cy5 incubated in mouse serum.

Fig. S5. Flow cytometry analysis of A20, untransduced CD8+ T, CAR T and Naïve T cells.

Fig. S6. GzmB activity under acidic condition.

Fig. S7. Bioluminescent assay confirmed the cytotoxic function of CD19-28ζ CAR T cells in the presence of G-SNAT or G-SNAT-Cy5.

Fig. S8. Cell viability study of G-SNAT and G-SNAT-Cy5.

Fig. S9. Confocal imaging of B16-mCD19 treated with activated untransduced CD8+ or CD19-28ζ CAR T cells.

Fig. S10. Illustration of the post-click labeling of G-SNAT and C-SNAT4 by Cy5 azide.

Fig. S11. Imaging gzmB and caspase 3 activity in GD2-4-1BBζ CAR T treated SB28-RFP/GD2 cells with G-SNAT-Cy5 and C-SNAT4-Cy5.

Fig. S12. Bioluminescence imaging and quantification of A20 implanted, activated untransduced CD8+ or CD19-28ζ CAR T^Luc+^ cells treated tumor-bearing mice with D-luciferin.

Fig. S13. Longitudinal fluorescence imaging of A20 implanted, activated untransduced CD8+ or CD19-28ζ CAR T^Luc+^ cells treated tumor-bearing mice with G-SNAT-Cy5.

Fig. S14. Longitudinal bioluminescence imaging of A20^Fluc+^ implanted, activated untransduced CD8+ or CD19-28ζ CAR T cells treated tumor-bearing mice with D-luciferin.

Fig. S15. Illustration of the gzmB activated bioluminescent assay with GBLI2.

Fig. S16. Longitudinal bright field imaging of nontreated, treated non-responded and responded mice.

Fig. S17. Relative fluorescent or bioluminescent intensity of tumors imaged with G-SNAT-Cy5, D-luciferin and GBLI2.

Fig. S18. Immunofluorescent staining analysis of nontreated, treated nonresponded and responded tumors.

Movie S1. Imaging of G-SNAT-Cy5 nanoaggregates in a single naïve CD8+ T cell.

Movie S2. Imaging of G-SNAT-Cy5 nanoaggregates in a single activated untransduced CD8+ T cell.

Movie S3. Imaging of G-SNAT-Cy5 nanoaggregates in a single CD19-28ζ CAR T cell.

Appendix S1. ^1^H and ^13^C nuclear magnetic resonance spectra.

## Supporting Information

### Materials and Methods

#### 1. General information – chemistry

Mouse gzmB substrate IEFD (Iso-Glu-Phe-Asp) with N-terminal acetylation was customized from GenScript USA (Piscataway, NJ). All chemicals were obtained from commercial sources unless otherwise stated. Reactions were monitored by thin layer chromatography (TLC) on 0.25 mm silica gel 60F plates. The ^1^H and ^13^C NMR spectra were acquired on Inova 500 MHz nuclear magnetic resonance (NMR) spectrometers at Stanford University. Data for ^1^H NMR spectra are reported as follows: chemical shifts are reported as δ in units of parts per million (ppm). Multiplicities are reported as follows: s (singlet), d (doublet), t (triplet), q (quartet), dd (doublet of doublets), quint (quintet), m (multiplet), or br (broadened); coupling constants are reported as a J value in Hertz (Hz); the number of protons (n) for a given resonance is indicated nH, and based on the spectral integration values. HRMS samples were performed on ESI-MS at the Waters Acquity UPLC and Thermo Exactive Orbitrap mass spectrometer at the Vincent Coates Foundation Mass Spectrometry Laboratory, Stanford University Mass Spectrometry. High-performance liquid chromatography (HPLC) was performed on a Dionex Ultimate 300 HPLC System (Thermo Scientific) equipped with a GP50 gradient pump and an in-line diode array UV-Vis detector. Reverse-phase C18 (Phenomenex, 5 μm, 4.6 × 250 mm or Dionex, 5 μm, 21.2 × 250 mm) columns were used with acetonitrile/water gradient mobile phase containing 0.1% trifluoroacetic acid (at a flow rate of 1 or 12 mL/min for analysis or purification respectively). TEM was performed on a JEOL JEM1400 transmission electron microscope. DLS and Zeta potentials were measured on Malvern ZetaSizer.

#### 2. Compound syntheses and characterizations

Synthesis of G-SNAT precursor is shown in Fig. S1.

Synthesis of G-SNAT and G-SNAT-Cy5 is shown in Fig. S2.

**(4*S*,10*R*,13*S*,16*S*,19*S*)-19-((2*S*,3*S*)-2-acetamido-3-methylpentanamido)-16-benzyl-13 (carboxymethyl)-1-(4-((3-((2-cyanopyrimidin-5-yl)oxy)propyl)carbamoyl)phenyl)-10-((ethylsulfinothioyl)methyl)-3,6,9,12,15,18-hexaoxo-4-(prop-2-yn-1-yl)-2,5,8,11,14,17-hexaazadocosan-22-oic acid (G-SNAT)**

i. To a mixture of Ac-IEFD-OH (0.05 mmol, 33.8 mg), HBTU (0.75 mmol, 28.4 mg) and compound **6** (0.05 mmol, 31.3 mg) was added DMF (15 mL), followed by *N*,*N*-Diisopropylethylamine (DIPEA, 0.25 mmol, 32.4 mg). The resulting solution was stirred for 2.5 h at r.t., then concentrated under high vacuum to remove DMF. The resulting residue was washed with water and brine, dried over anhydrous Na_2_SO_4_, filtered, and concentrated.
ii. The remained residue was added with a solution of CF_3_COOH : DCM : TIPS = 1 : 1 : 0.05 (10 mL) and continued to stir at r.t. for another 2 h. After concentration, the final product **G-SNAT** was purified by preparative-HPLC to afford a white powder (10.0 mg, 17.0%). ^1^H NMR (500 MHz, DMSO-*d^6^*) δ 9.10 (s, 1H), 8.60 (t, *J* = 6.0 Hz, 1H), 8.50 (t, *J* = 5.7 Hz, 1H), 8.39 (d, *J* = 7.7 Hz, 1H), 8.18 – 8.12 (m, 3H), 7.94 (dd, *J* = 17.9, 8.0 Hz, 2H), 7.85 (d, *J* = 8.0 Hz, 1H), 7.77 (d, *J* = 8.2 Hz, 2H), 7.33 (d, *J* = 8.1 Hz, 2H), 7.22 (d, *J* = 4.3 Hz, 4H), 7.15 (dt, *J* = 8.7, 4.2 Hz, 1H), 4.60 (q, *J* = 7.2 Hz, 1H), 4.53 (d, *J* = 8.6 Hz, 1H), 4.51 – 4.45 (m, 2H), 4.45 (d, *J* = 6.7 Hz, 2H), 4.38 (dd, *J* = 15.7, 6.1 Hz, 1H), 4.30 (dd, *J* = 15.8, 5.8 Hz, 1H), 4.19 (q, *J* = 7.9 Hz, 1H), 4.10 (t, *J* = 7.8 Hz, 1H), 3.77 (qd, *J* = 16.7, 5.7 Hz, 2H), 3.11 (dd, *J* = 13.5, 4.8 Hz, 1H), 3.03 (dd, *J* = 14.0, 4.0 Hz, 1H), 2.94 – 2.85 (m, 2H), 2.82 – 2.65 (m, 4H), 2.64 – 2.56 (m, 1H), 2.53 (d, *J* = 7.4 Hz, 2H), 2.22 – 1.97 (m, 4H), 1.85 (s, 3H), 1.81 (s, 1H), 1.68 (dd, *J* = 15.6, 8.5 Hz, 2H), 1.38 (s, 1H), 1.21 (q, *J* = 6.5 Hz, 3H), 1.06 (dt, *J* = 14.7, 7.6 Hz, 1H), 0.82 – 0.73 (m, 6H). ^13^C NMR (126 MHz, DMSO-*d^6^*) δ 174.0, 171.8, 171.3, 171.1, 170.9, 170.7, 170.0, 169.7, 169.5, 168.5, 166.1, 165.1, 163.4, 142.2, 137.6, 133.0, 129.1, 128.0, 127.1, 126.8, 126.2, 115.6, 102.2, 80.5, 73.2, 66.4, 57.1, 53.6, 52.4, 51.9, 51.7, 42.2, 41.9, 40.5, 40.0, 39.9, 39.9, 39.8, 39.7, 39.5, 39.4, 39.2, 39.0, 37.6, 36.2, 36.0, 31.7, 30.1, 28.4, 27.2, 24.5, 22.5, 21.8, 15.4, 14.3, 11.0. MS: m/z calcd for C_54_H_68_N_12_O_14_S_2_ 1172.4; found 1171.3, [M + H]^+^

**1-(6-((3-(4-((2*S*,8*R*,11*S*,14*S*,17*S*,20*S*)-14-benzyl-20-((*S*)-sec-butyl)-17-(2-carboxyethyl)-11-(carboxymethyl)-2-((4-((3-((2-cyanopyrimidin-5 yl)oxy)propyl)carbamoyl)benzyl)carbamoyl)-8-((ethylsulfinothioyl)methyl)-4,7,10,13,16,19,22-heptaoxo-3,6,9,12,15,18,21-heptaazatricosyl)-1H-1,2,3-triazol-1-yl)propyl)amino)-6-oxohexyl)-3,3-dimethyl-2-((1E,3E)-5-((E)-1,3,3-trimethyl-5-sulfonatoindolin-2-ylidene)penta-1,3-dien-1-yl)-3H-indol-1-ium-5-sulfonate (G-SNAT-Cy5)**

Compound G-SNAT (6.1 µmol, 7.1 mg) and sulfo-Cy5-azide (6.5 µmol, 5.0 mg) was dissolved in DMSO (200 µL) and 0.1 M HEPES solution (800 µL), then CuSO_4_ (100 µL of 0.1 M stock solution in H_2_O), (BimC_4_A)_3_ (100 µL of 30 mM stock solution in H_2_O) and sodium ascorbate (100 µL of 1 M stock solution in H_2_O, freshly prepared) were added to the mixture. After stirring at r.t. for 1 h, the mixture was directly purified by preparative-HPLC to final product **G-SNAT-Cy5** as a blue powder (3.2 mg, 28.1%). MS: m/z calcd for C_89_H_111_N_18_O_21_S_4_ 1895.7; found 947.8, [M + 2H]^2+^; HRMS (ESI/Q-TOF): [M + 2H]^2+^ m/z calcd for C_89_H_111_N_18_O_21_S_4_ 949.3639; found 949.3642.

#### 3. General information – biology

Recombinant mouse granzyme B protein (C-6His, C765) was purchased from Novoprotein Scientific (Summit, NJ). Recombinant mouse active cathepsin C/DPPI Protein (2336CY010) was purchased from R&D systems (Minneapolis, MN). Recombinant human caspase-3 protein (235417) was purchased from Sigma-Aldrich (St Louis, MO). Granzyme B inhibitor (368055) was purchased from EMD Millipore (Burlington, MA). Rat anti-granzyme B mouse monoclonal antibody (16G6) was from Invitrogen (Carlsbad, CA). Anti-mouse PD-1 (clone-RMP1-14) and anti-mouse CTLA-4 (clone-9D9) were purchased from BioXcell (Lebanon, NH). IRDye 680RD goat anti-rat IgG, IRDye 680RD donkey anti-mouse IgG and IRDye 800CW donkey anti-rabbit secondary antibodies were purchased from LI-COR Biosciences (Lincoln, NE). FITC conjugated anti-mouse CD3 (17A2) and Alexa Fluor 594 anti-mouse CD19 (6D5) antibodies were purchased from Biolegend (San Diego, CA). Rabbit anti-β Actin polyclonal antibody (PA1-46296) was from Invitrogen. D-luciferin, potassium salt was purchased from Gold Biotechnology (St Louis, MO). Cyanine 5 azide was purchased from Lumiprobe (Hunt Valley, Maryland). RIPA buffer, Pierce BCA protein assay reagents and NuPAGE 4-12% Bis-Tris protein gels were purchased from Thermo Fisher Scientific (Waltham, MA). Microtube pestles were purchased from USA Scientific (Ocala, FL). Mouse serum was purchased from Sigma-Aldrich (St Louis, MO). Hoechst 33342 was purchased from Sigma-Aldrich. High-precision cover glass 22 × 22 mm with thickness 170 μm ± 5 μm (0107052) was purchased from Marienfeld-Superior (Lauda-Königshofen, Germany). Superfrost Plus microscope slides (12-550-15) were purchased from Fisher Scientific. SlowFade Diamond antifade mountant (S36963) was purchased from Invitrogen. Western blots were imaged using a LI-COR Odyssey imaging system.

#### 4. Cell culture

A20 (TIB-208) murine B lymphocyte cell line was obtained from ATCC and cultured in RPMI 1640 medium supplemented with 10% fetal bovine serum (FBS), 100 U/ml penicillin, 100 ug/ml streptomycin, 2 mM L-glutamine, 0.05 mM 2-mercaptoethanol, 10 mM HEPES, 1 mM sodium pyruvate, 4500 mg/L glucose, and 1500 mg/L sodium bicarbonate. Culture was maintained in 37°C with 5% CO_2_. The modified A20 FLuc+/YFP/neo was generated as previously reported^1^. Cells were grown in RPMI-1640 medium (Sigma-Aldrich) supplemented with 10% fetal bovine serum (ThermoScientific), 2 mM L-glutamine (Sigma) and 0.05 mM 2-mercaptoethanol (Sigma-Aldrich, St. Louis, MO) at 37 °C in 5% CO_2_ atmosphere. B16 melanoma cell line was obtained from ATCC. SB28, a murine glioblastoma cell line was generously provided from Dr. Hideho Okada (University of California San Francisco, San Francisco, CA). Both cell lines were maintained in DMEM supplemented with 10% fetal bovine serum (FBS) and 1% antibacterial/antimycotic solution (Catalog#15240112, ThermoFisher). B16 and SB28 lines were engineered to express murine CD19-TurboRFP-Rluc and GD2-TurboRFP, respectively, using lentiviral transduction followed by three rounds of cell sorting for the highest 2.5% of TurboRFP expressers. CT26 murine colon carcinoma cell line was obtained from ATCC and cultured in RPMI 1640 medium supplemented with 10% fetal bovine serum (FBS), 100 U/ml penicillin, 100 ug/ml streptomycin, and 2 mM L-glutamine. Culture was maintained in 37°C with 5% CO_2_.

All cell lines were routinely tested for mycoplasma contamination (MycoAlert Mycoplasma Detection Kit purchased from Lonza), with authentication performed at Stanford Functional Genomics Facility for Short Tandem Repeat (STR) profiling.

#### 5. Viral vectors

MSGV retroviral vectors encoding second generation CARs, CD19-28ζ or GD2-4-1BBζ CAR^2^, were used for construction of CD19(murine)-targeted CART cells or GD2-targeted CAR T cells. 293GP producer cell lines, provided as a gift from Dr. Crystal Mackall (Stanford University, Stanford, CA), produced retroviral supernatant harboring CD19-28ζ or GD2-4-1BBζ CAR vectors. Supernatant was collected after 72 hours of 293GP culture, centrifuged to discard cell debris, and stored at -80°C for 6 months.

For retroviral transduction, non-tissue culture treated 6-well plates (Catalog# 3736, Corning, NY) were coated with Retronectin (Takara, Japan) diluted in PBS (24 µg/ml) for 16 hours at 4°C. Plates were blocked with 2% BSA in PBS for 30 minutes and then discarded. Frozen supernatants were thawed on ice and 2 ml supernatant was mixed with 1 ml T cell culture medium and added to each well, followed by centrifugation for 3 hours at 3200 rpm at 32°C. Supernatants were discarded and plates were ready for T cell addition.

#### 6. Generation of CD19-28ζ and GD2-4-1BBζ CAR T cells

Splenocytes were harvested from 8–10 weeks old, female BALB/c mice (Charles River Laboratories, Wilmington, MA) or 6–8 weeks old, female C57Bl/6J mice (Jackson Laboratory, Bar Harbor, ME) and processed in 1× PBS (Invitrogen) supplemented with 2% FBS (Invitrogen) into single-cell suspensions. CD8+ T cells were prepared using EasySep™ Mouse CD8+ T Cell Isolation Kit (Catalog# 19853, Stemcell Technologies, Vancouver, Canada). Total T cells were prepared using EasySep™ Mouse Total T Cell Isolation Kit (Catalog# 19851, Stemcell Technologies, Vancouver, Canada), as recommended by the manufacturer. In brief, cells were blocked with normal rat serum, stained with Isolation cocktail antibodies, mixed with Streptavidin RapidSpheres, and purified cells were isolated manually using MACS EasySep™ magnetic column (Catalog #18000, Stemcell Technologies, Vancouver, Canada. This purification protocol yielded >90% purity.

CD19-28ζ and GD2-4-1BBζ CAR-T cells were genetically engineered from CD8+ T cells and total T cells, respectively, using retroviral transduction^3^. The enriched T cell fractions were activated with magnetic beads coated with agonistic anti-CD3 and CD28 antibodies (Catalog# 11456D, ThermoFisher, Waltham, MA) at 1 : 1 (beads : cells) ratio in T cell medium (RPMI-1640 medium supplemented with 10% fetal bovine serum, 2 mM L-glutamine, 1% antibacterial/antimycotic solution, 10 U/ml recombinant murine IL-2 (Peprotech, NJ), 10 U/ml recombinant murine IL-7 (Peprotech, NJ), and 0.05 mM 2-mercaptoethanol). After 24 hours, 1 × 10^6^ activated T cells were transduced with retroviruses bound to retronectin-coated tissue culture plates for 3 days, after which beads were removed. Transduced and untransduced T cells were maintained at 1×10^6^ cells per milliliter in T cell medium and analyzed using flow cytometric analysis. For mice studies, T cells were transferred three days after removal of beads and viral supernatant. CAR T cells generated from BALB/c mice were used to challenge A20 cells, while those produced from C57Bl/6J were used to challenge B16 cells.

#### 7. Flow cytometry

For T cell sorting, cells were harvested, washed twice with 1× PBS, and re-suspended in cold PBS containing 2% FBS (at a density of 1 × 10^6^ cells/100µl). Subsequently, primary labelled antibodies or Pierce™ Recombinant Protein L (catalog #21189, ThermoFisher) were added into the cell suspension according to the manufacturer’s instructions and incubated for 20 minutes at 4 °C in the dark. Immediately after the incubation, the cells were washed thrice with ice cold PBS. For the protein L treated cells, APC-labelled streptavidin was added to cells and incubated for 15 minutes at 4 °C in the dark. Flow cytometry was performed using a BD LSRII and FlowJo Software (TreeStar) for analysis. Gating strategies for detecting the percentages of naïve, CD8+ T cells and CAR-T cells were analyzed using the following antibodies: anti-CD3 (clone:17A2, Brilliant Violet 421), anti-CD8 (clone:53-6.7, FITC or clone:YTS156.7.7, PE/Dazzle 594), anti-CD44 (clone: IM7, PE), anti-CD19 (PE/APC/FITC), anti-CD62L (clone: MEL-14, Alexa Fluor 700), anti-CTLA4 (clone UC10-4B9, PE), anti-PD1 (clone: 29F.1A12, APC/Fire 750), and streptavidin (APC). Dead cells were excluded by using Zombie NIR™ Fixable Viability kit (catalog# 423105). The antibodies used for flow cytometric analysis were obtained from Biolegend.

For analysis of CAR or CD8+ T cells treated A20 cells as suspension in the presence of G-SNAT-Cy5, cells were centrifuged gently at 300 g for 10 min, washed with cold PBS once, then analyzed with a 4-laser, 12-color DxP12 Cytek upgrade (Becton Dickinson, Cytek Biosciences) and evaluated according to their size (FSC, Front Scatter) on a linear scale. For Cy5, cells were excited by 640 nm later, filtered by 655 nm, LP (Long Pass filter) and 661/716 nm, BP (Band Pass filter). For FITC α-CD3 antibody, cells were excited by 488 nm later, filtered by 560 nm, SP (Short Pass filter) and 525/550 nm, BP. For Alexa Fluor 594 conjugated α-CD19 antibody, cells were excited by 561 nm later, filtered by 600 nm, SP (Short Pass filter) and 590/620 nm, BP. Flow cytometry data analysis and 3D dimensional projection was done using FlowJo V10 software. The MFI (mean fluorescence intensity) was collected and plotted.

#### 8. Western blot analysis of cells and tumors

Cells were trypsinized, pelleted and washed 3 times with ice cold PBS and lysed in RIPA buffer. Tumor tissues were cut into small pieces and grinded in 1.5 mL Eppendorf tubes with microtube pestles for lysis in RIPA buffer. Protein concentration of centrifuged whole lysate were determined with BCA assay. Tumor lysates were loaded in NuPAGE 4-12% Bis-Tris protein gels for electrophoresis at 200V for 90 min. Wet transfer were performed using a Bio-Rad transfer kit at 300 mA for 90 min. The transferred nitrocellulose membrane was blocked in PBS containing 5% BSA and 0.1% Tween-20 for 1 h. Primary antibody incubation was performed in the blocking buffer overnight at 4 °C in recommended concentration. The membrane was then washed with PBS containing 0.1% Tween-20 for 5 min, four times. IRdye conjugated secondary antibody incubation was performed in the blocking buffer for 2 h at room temperature. After washing four times with PBS containing 0.1% Tween-20, membranes were analyzed in a LI-COR Odyssey imaging system.

#### 9. Granzyme B-mediated macrocyclization *in vitro*

G-SNAT (10 μM, 10 μL of 1 mM stock in DMSO) was diluted in reaction buffer containing 50 mM Tris at pH 7.5 with or without 1 mM TCEP (2 μL of 500 mM stock in water) and 1 μg recombinant mouse gzm B in 1 mL final concentration. Reactions were performed at 37 °C overnight and monitored by HPLC. For kinetic study, G-SNAT (10 μM) was incubated with equal amount (100 U) of recombinant mouse GzmB (0.05 μg/ml) in Tris buffer and human caspase-3 in a buffer containing 50mM HEPES, 100 mM NaCl, 1 mM EDTA and 10% glycerol for up to 8 hours.

#### 10. Quantitative real time PCR

Total RNA was extracted from 1 million of each T cell population tested using RNeasy Mini kit (Qiagen, Hilden, Germany) following the manufacturer’s instructions. cDNA synthesis was conducted using iScript cDNA synthesis kit (Bio-Rad, Hercules, CA) as follows: 1 µg sample RNA mixed with 4 µL of 5x iScript reaction mix and 1µL of iScript reverse transcriptase in total volume of 20 µL. Reaction protocol as follows: 5 min at 25 ^°^C for priming, 20 min at 46 ^°^C for reverse transcription, 1 min at 95 ^°^C for inactivation of reverse transcription. PrimePCR™ FAM-conjugated Probe Assays (Bio-Rad, Hercules, CA) for murine granzyme B (Assay ID: qMmuCEP0053372) and GAPDH (Assay ID: qMmuCEP0039581) were used with Sso Advanced Universal Probes Supermix (Bio-Rad) to run quantitative PCR reactions as follows: 100 ng of cDNA mixed with 10 µl of Sso Advanced universal probes supermix(2x) and 1 µl of gene-specific hydrolysis probe in final volume of 20 µl. Reaction protocol for qPCR as follows: 30 seconds at 95 ^°^C for cDNA denaturation, followed by 40 cycles of 15 seconds at 95 ^°^C for denaturation, 30 seconds at 60 ^°^C for annealing/extension. All cDNA synthesis and qPCR reactions were conducted using a CFX96 Real-Time System C1000 Touch Thermal Cycler (Bio-Rad). Control reactions without DNA template or reverse transcriptase did not show amplification. Technical replicates for all samples were performed in triplicates. Relative gene expression was measured using the ΔΔCT method.

#### 11. Post-click labeling of G-SNAT by Cy5-azide for epifluorescence microscope imaging

For imaging with microscope, SB28 cells were seeded at about 50% confluence on cover glasses in a 6-well plate with complete medium a day before. After incubation at 37 °C for 3.5 h with 20 μM G-SNAT (from 10 mM stock in DMSO) and GD2-4-1BBζ CAR-T cells at 1 : 2 ratio in medium containing 1 % FBS, cells were washed 3 times with HBSS then fixed with 10% formalin for 30 min. Cells were permeabilized with PBS-Triton X100 (0.1% v/v) at room temperature for 15 min then washed 3 times with PBS. Post-click assay solution was freshly prepared with 100 mM ascorbic acid (100 μL of 1 M stock in water, freshly prepared), 1 mM CuSO_4_ (10 μL of 100 mM stock in water), 15 μM (BimC_4_A)_3_ (0.5 μL of 30 mM stock in water) and 5 μM Cy5-azide (1 μL of 5 mM in DMF) in 1 mL PBS. Cells were incubated with assay solution for 4 h at 37 °C then washed 3 times with PBS. Cells were then stained with 300 nM DAPI in PBS for 10 min at room temperature, washed with PBS 3 times and mounted on glass slides with antifade mounting medium.

The epifluorescence microscope images were acquired by 1X81 inverted microscope (Olympus) equipped with pE-4000 illumination systems (CoolLED) and ORCA-Flash4.0 digital CMOS camera (HAMAMATSU) with excitation at 405 nm for DAPI and 650 nm for Cy5. Digital images were reconstructed by MetaMorph software (v. 7.8.11.0) and analysed using the ImageJ (NIH) software package.

#### 12. Confocal microscope imaging and flow cytometry analysis

Confocal images were obtained by a Zeiss LSM710 inverted confocal microscope, using a 20x, 40x/oil or 63x/oil immersion objective. All fluorescence images were gathered sequentially and stacked. Sequential Z sections of stained cells were recorded for generation of stacked images. Multi-channel 3D projections of fluorescent images were constructed from sequential Z sections of cells assembled in ImageJ.

#### 13. Serum stability

G-SNAT-Cy5 (100 μM) was diluted in 100 μL mouse serum and incubated at 37 °C for 0, 30 min, 1, 2, 4, 6 and 8 h. Serum proteins were denatured by mixing cold methanol (900 μL) and precipitated by centrifugation at 8,000 g for 10 min. Supernatant were analyzed by HPLC (fluorescent detector, Ex640/Em670) and LC-MS. Percentage of tracer (relative area) was calculated as (peak area of tracer/total peak area on the HPLC chromatogram) × 100.

#### 14. CAR T cell cytotoxic function study and cell viability assay

For cytotoxic function, 2.5 × 10^5^ A20^Luc+^ cells were incubated without or with effector CD19-28ζ CAR T cells at 1:10 and 1:1 effector-to-target (E:T) ratios in the presence of G-SNAT or G-SNAT-Cy5 at 2.5 μM, 5 μM. or solvent control (DMSO, 1%) overnight in a 96-well black plate. The final volume was 200 μL. Next day, D-luciferin was added to 300 μg/mL final concentration before incubation at 37 °C for 5 min and scanned with an IVIS optical imager. The bioluminescence intensity was collected by defining the ROI on each well. Percent survival was calculated over the bioluminescent intensity of A20^Luc+^ cells without CAR T cells in each treatment group.

Cell viability assay was performed with a CellTiter 96 AQueous One Solution Cell Proliferation Assay (MTS) kit (Promega, Madison, WI) according to the manufacturer’s protocol. Briefly, 2.5 × 10^5^ A20^Luc+^, 2 × 10^5^ activated untransduced CD8+ and 2 × 10^5^ CAR T cells (70.7% transduction efficiency, so total 2.83 × 10^5^ CAR T and CD8+ T cells) were incubated with G-SNAT (5 μM – 100 μM), G-SNAT-Cy5 (5 μM – 100 μM), and solvent (DMSO, 1%) in a 96 well-plate at 37 °C for 3.5 h in the presence of MTS reagent. The absorbance at 490 nm was collected on a plate reader and plotted.

#### 15. *In vivo* imaging and data analysis

All experimental procedures using mice were performed in agreement with protocols approved by Institutional Animal Care and Use Committee (IACUC) of Stanford University as well as the USAMRMC Animal Care and Use Review (ACURO) in accordance with the laws of the United States and regulations of the Department of Agriculture. BALB/c and BALB/c Rag2^−/−^γc^−/−^ mice were purchased from Jackson Laboratory and bred in house at the Research Animal Facility at Clark Center of the Stanford University (Stanford, CA).

For the A20 subcutaneous model, 1 × 10^6^ A20 cells were injected subcutaneously into right upper flanks of 6-week-old female BALB/c Rag2^−/−^γc^−/−^ mice in 90% Matrigel. Tumor burden was assessed by external caliper and calculated by use of the formula (Length × Width × height)/2. Tumors were grown to around 300 mm^3^ for imaging. 6 × 10^6^ CD19-28ζ CAR T cells prepared from immunocompetent Fluc+ BALB/c mice were intratumorally injected in a volume of 200 μL in PBS. To adjust for the variable transduction efficiencies of CD19-28ζ retroviral vectors into CD8+T cells, the concentrations of injected CD19-28ζ CAR T cells were normalized.

For the systemic lymphoma model, BALB/c mice were sublethally (4.4 Gy) irradiated. 1 × 10^6^ A20 luc+/YFP cells were injected via tail vein. 2.4 × 10^6^ CAR-T or CD8+T cells prepared from immunocompetent BALB/c mice were injected through the retro-orbital route. To adjust for the variable transduction efficiencies of CD19-28ζ retroviral vectors into CD8+T cells, the concentrations of injected CD19-28ζ CAR T cells were normalized. Tumor burden was assessed by *in vivo* BLI^4^. Briefly, 10 min after intraperitoneal injection of D-luciferin, mice were imaged for 5 min with an IVIS 100 charge-coupled device imaging system (Xenogen). For G-SNAT-Cy5, 5 nmol probes were injected through tail vein and imaged longitudinally with IVIS (Ex640/Em690) from 1 to 20 h. Images were analyzed with Living Image Software 2.5 (Xenogen). Intensity was collected by defining region of interest (ROI).

For the checkpoint blockade therapy model, 1 × 10^6^ CT26 cells were injected subcutaneously into right upper flanks of 8-week-old female Fluc+ BALB/c mice in 50% Matrigel. Tumor burden was assessed by external caliper and calculated by use of the formula (Length × Width × height)/2. When reached ∼200 mm^3^, tumors were treated with a combined regimen of anti-PD1 (200 μg) and anti-CTLA4 (100 μg) intraperitoneally for three rounds at day 9, 12, and 15. Treated mice with complete tumor regression were defined as responders. The mice were imaged sequentially with GBLI-2 (200 μg-i.v. retro orbital), G-SNAT-Cy5 (5 nmol-i.v. retro orbital, Ex), and D-luciferin (3 mg-i.p.), two hours apart, at day 9, 12, 15, 18, and 20 with a Largo-X imaging system. Images were analyzed with Aura image software 3.2 (Spectral Instruments Imaging). Intensity was collected by defining ROI on tumors.

#### 16. Immunofluorescent staining

Tissue preparation: The animals were anesthetized and sacrificed by a cervical dislocation after CO_2_ narcosis. Tumors were surgically removed post-mortem and fixed overnight in sterile 4% paraformaldehyde and transferred in 30% sucrose for 24 h. After removal of PFA by sucrose treatment, the tissues were cut to desired size and embedded in peel-a-way disposable base molds (ThermoFisher Scientific) containing O.C.T (Tissue-Tek O.C.T. Compound, Sakura Finetek) and frozen using dry ice. The embedded frozen tissues were cut to 10-micron thickness using CryotomeTM and the tissue slices were collected on Superfrost Ultra Plus glass slides (ThemoFisher Scientific). The glass slides that contained the tissue slices were stored at -20°C.

Histology: The tumor tissue slides were equilibrated to room temperature before any further manipulation. The slides were immersed in antigen retrieval solution containing 10 mM citric acid (monohydrate) and 0.05% Tween 20 at pH 6.0 and heated to 98 °C for 20 min. After elapse of 20 min, the slides were removed gently and cool-downed to room temperature, allowing reformation of antigenic sites after exposure to high temperature. The slides were washed twice using wash buffer (1x Tris Buffer Saline (TBS) plus 0.025% Triton X-100) with gentle shaking in orbital shaker. The tissues were then blocked using 10% goat serum in blocking buffer (1x TBS containing 1% BSA) for 2 h at room temperature, followed by overnight primary antibody treatment at 4 °C. The primary antibodies used were Rabbit anti-mouse Granzyme B (ab4059, Abcam at 1:100 dilution in blocking buffer) and Rat anti-mouse CD8a (MCA609GT, Bio-Rad at 1:100 dilution). After primary antibody staining, the tissues slides were washed twice and stained with secondary antibodies conjugated to fluorophore. The secondary antibodies used were Alexa Fluor 488-conjugated Goat anti-rabbit secondary (A11034, Life technologies at 1:200 dilution) and PerCP/Cy5.5-conjuagted Goat anti-rat IgG secondary (405424, Biolegend at 1:100 dilution). This step was followed by twice washing and staining with primary antibody for CD19 pre-conjugated with fluorophore, Alexa Fluor 594-conjugated Rat anti-mouse CD19 (115552, Biolegend at 1:100 dilution). The tissues slides were then stained with 300 nM DAPI solution for 20 min and air dried and mounted using ProLongTM Diamond anti-fade mountant (ThermoFisher Scientific) and imaged using a Zeiss LSM710 confocal microscope.

#### 17. Statistical Analysis

GraphPad Prism 7 was utilized for plotting and statistical analysis. The significant difference was determined by performing one-way (figure) or two-way (figure.) ANOVA followed by Bonferroni’s multiple comparisons test to determine the statistical significance with 95% confidence intervals with \**p* < 0.0332; \*\**p* < 0.0021, \*\*\**p* < 0.0002, \*\*\*\**p* < 0.0001, ns: not significant.

## SUPPLEMENTARY FIGURES

**Fig. S1.**
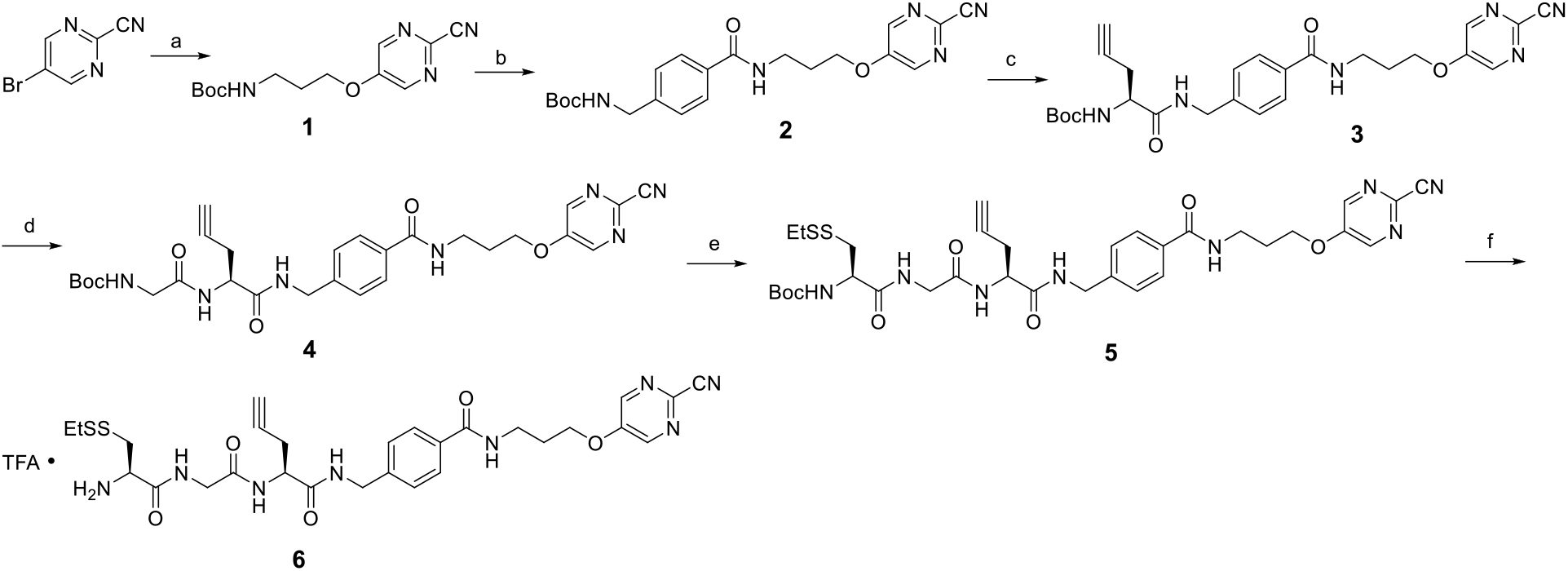
Synthesis of G-SNAT precursor. Reagents and conditions: (a) 3-(Boc-amino)-1-propanol, Pd(OAc)_2_, BINAP, Cs_2_CO_3_, reflux, 110 °C; (b) (i) 25% TFA in DCM, r.t., 30 min; (ii) 4-(Boc-aminomethyl)benzoic acid, HBTU, DIPEA, DMF, r.t., 2 h. (c) (i) 25% TFA in DCM, r.t., 30 min; (ii) Boc-propargyl-Gly-OH, HBTU, DIPEA, DMF, r.t., 2 h. (d) (i) 25% TFA in DCM, r.t., 30 min; (ii) Boc-Gly-OH, HBTU, DIPEA, DMF, r.t., 2 h. (e) (i) 25% TFA in DCM, r.t., 30 min; (ii) Boc Cys(SEt)-OH.DCHA , HBTU, DIPEA, DMF, r.t., 2 h; (f) 25% TFA in DCM, r.t., 30 min.

**Fig. S2.**
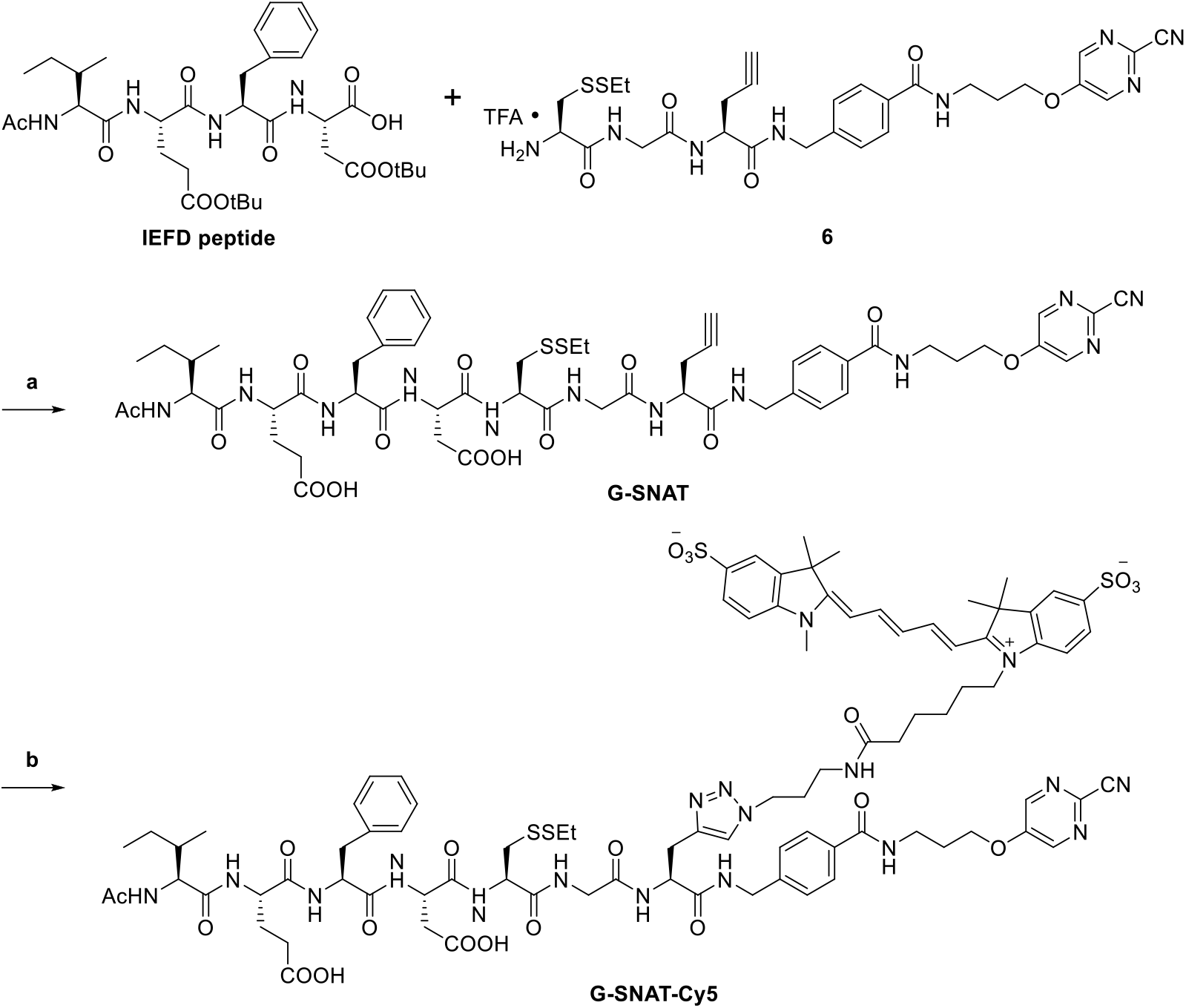
Synthesis of G-SNAT and G-SNAT-Cy5. Reagents and conditions: (a) (i) HBTU, DIPEA, DMF, r.t., 2.5 h; (ii) CF3COOH: DCM: TIPS = 1: 1: 0.05, 5 mL, r.t., 2 h; (b) sulfo-Cy5-azide, CuSO4, sodium absorbate, (BimC4A)3, DMSO/HEPES = 1: 4.

**Fig. S3.**
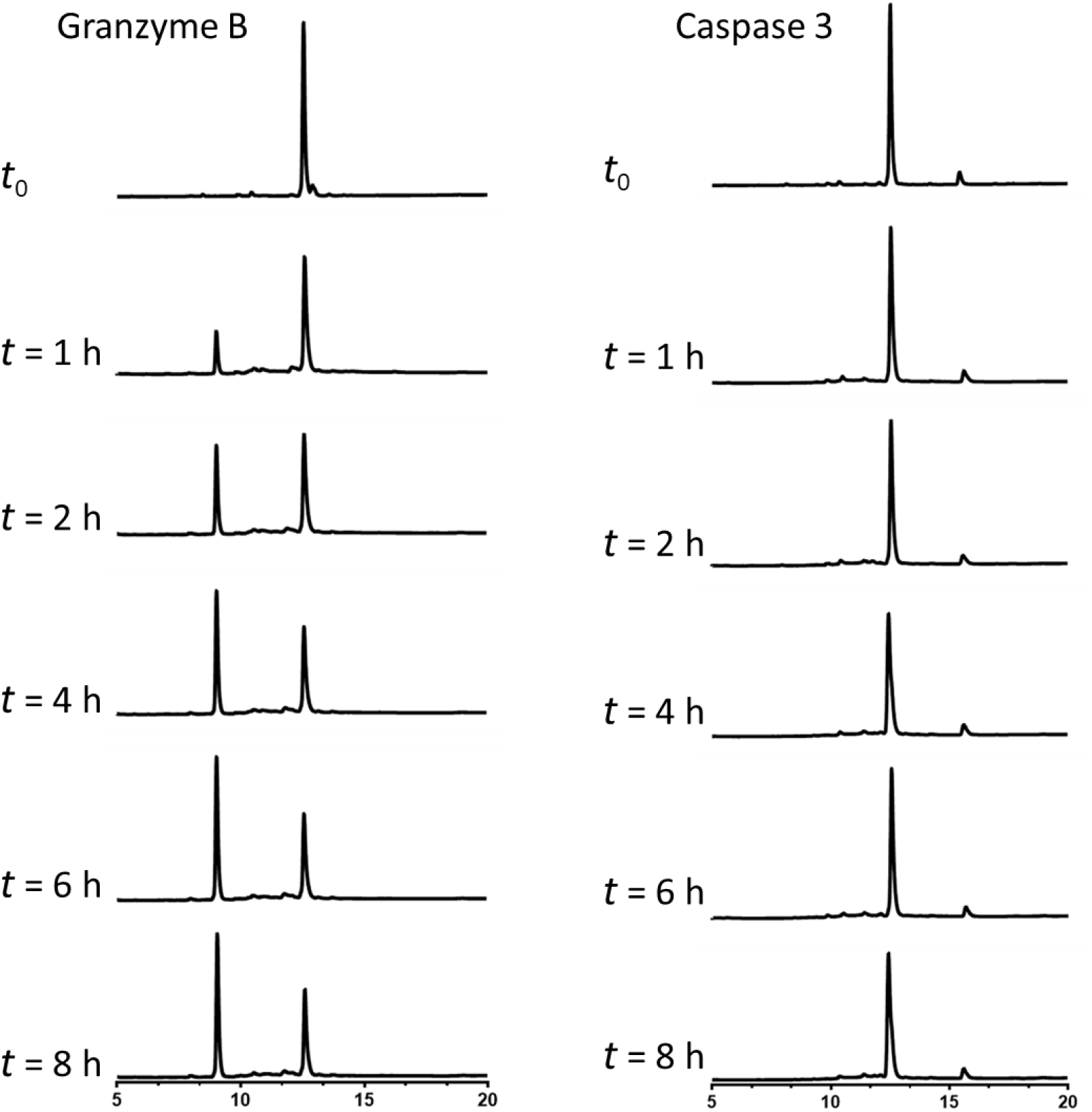
Representative HPLC traces in gzmB and caspase-3 kinetic study. The enzymatic reaction kinetics and specificity studies by longitudinal monitoring of percentage conversion of G-SNAT (10 μM) into cleaved G-SNAT after incubation with equal amounts (100 U) of recombinant mouse

**Fig. S4.**
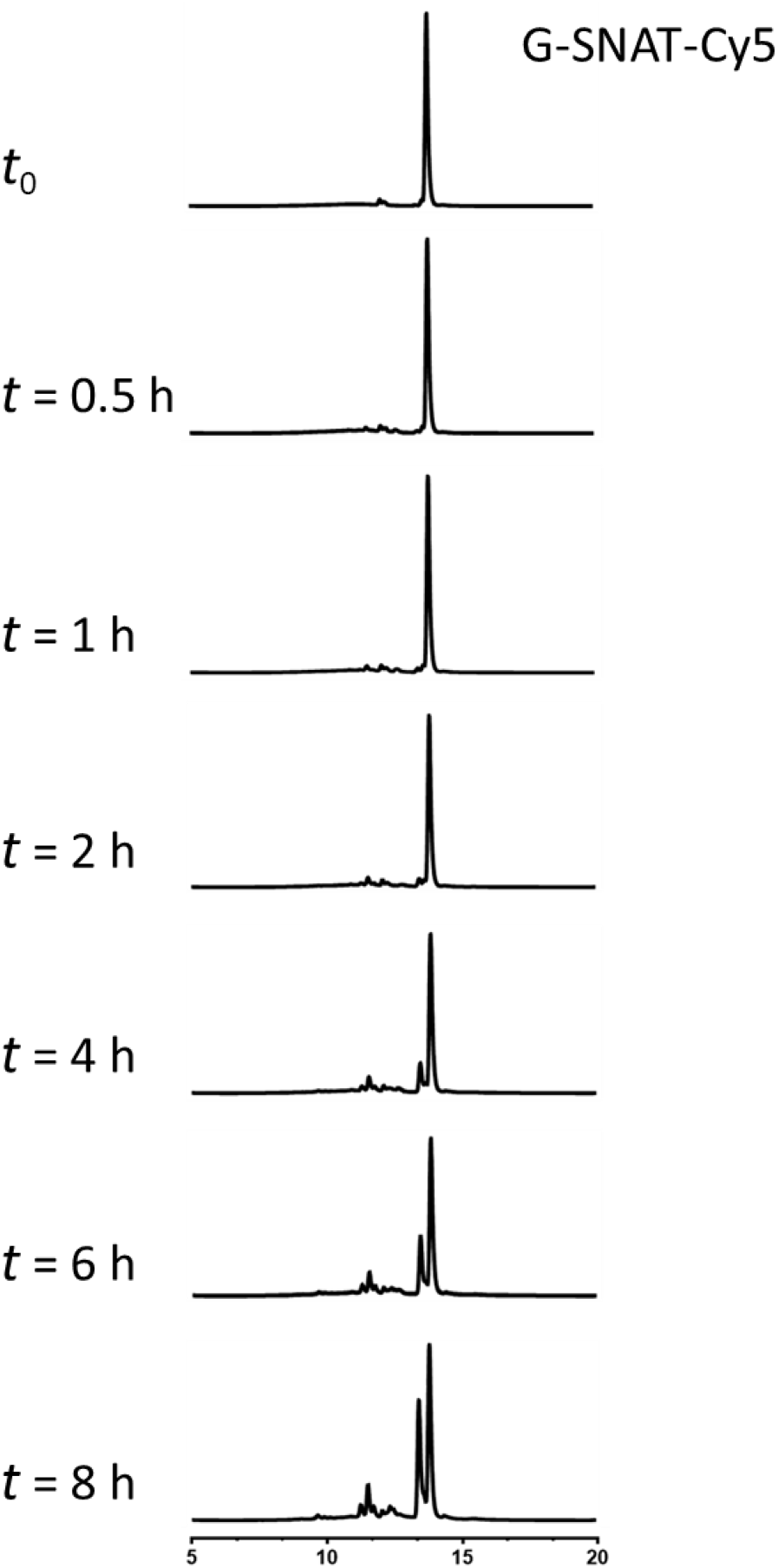
Representative HPLC traces of 100 μM of G-SNAT-Cy5 incubated in mouse serum at different times (fluorescent detector Ex640/Em670). Samples were dissolved and precipitated in 80% methanol before HPLC analysis.

**Fig. S5.**
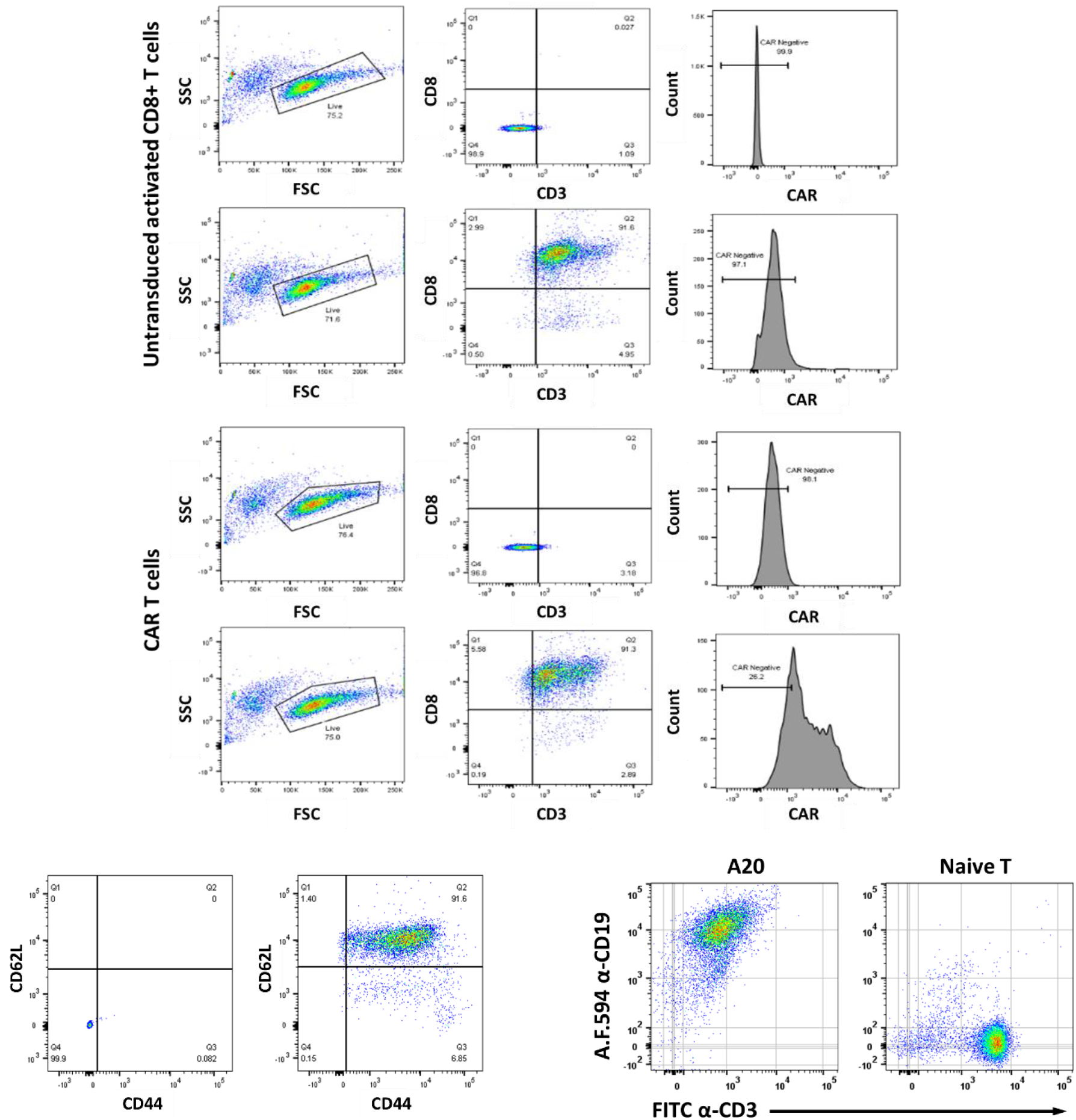
Flow cytometry analysis of A20, untransduced CD8+ T, CAR T and Naïve T cells. Upper panel: Gating strategy used for identification of untransduced CD8+ and CAR T cells. Live cells were first identified and selected based on CD3+ (BV421) and CD8+ (APC/Cy7) staining. Final identification of CAR T cells was based on CAR (APC) staining. Representative histograms for the indicated cells, including a negative control with isotype staining is shown for untransduced CD8+ and CAR T cells. Lower left: Gating strategy used for identification of naïve T cells isolated from mice spleens. Live cells were first identified and selected based on CD3+ (BV421) and CD8+ (APC/Cy7) staining. Final identification of naïve T cells was based on CD44^low^(PE) and CD62^high^(APC) staining. Representative histograms for the indicated cells, including a negative control with isotype staining (Left) and staining with CD44 and CD62L. Lower right: The expression of CD19 on A20 cells was confirmed by staining with Alexa Fluor 594 conjugated CD19 antibody with naïve T cells (CD3+CD19-) as a control.

**Fig. S6.**
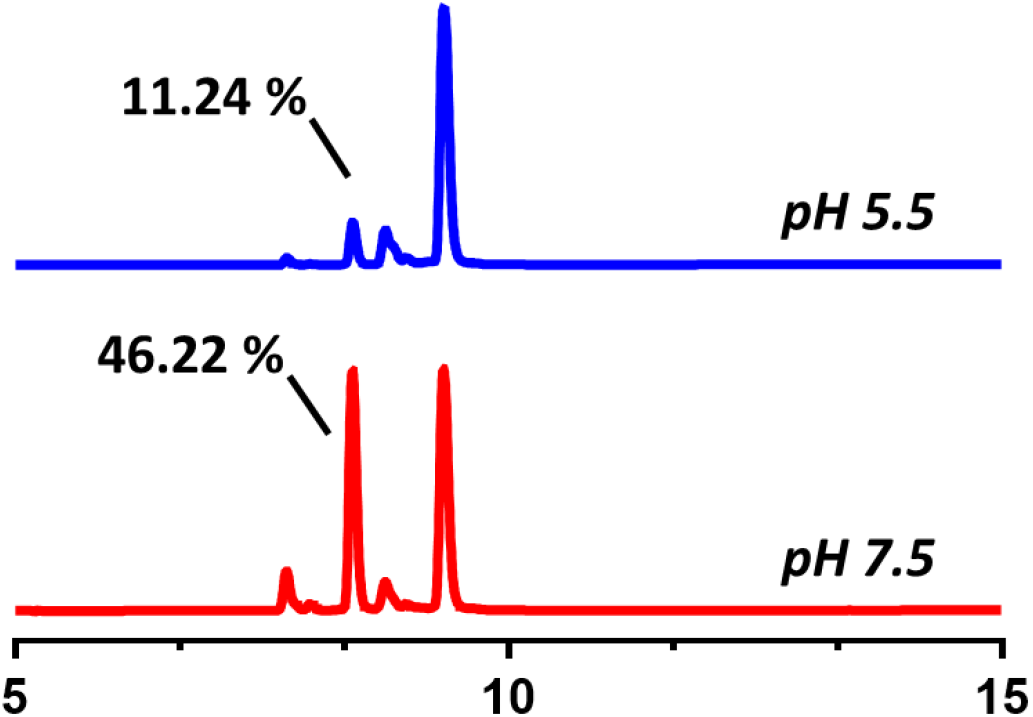
GzmB activity under acidic condition. HPLC traces of G-SNAT-Cy5 incubated with gzmB (0.05 μg/ml) in MES buffer (pH5.5, blue) or Tris buffer (pH7.5, red) at 37 °C for 4 hours. 11.24% and 46.22% indicate the percentage conversion relative to G-SNAT-Cy5 peak obtained by calculating the percentage of peak area (mAU*min) of probe on the corresponding HPLC trace.

**Fig. S7.**
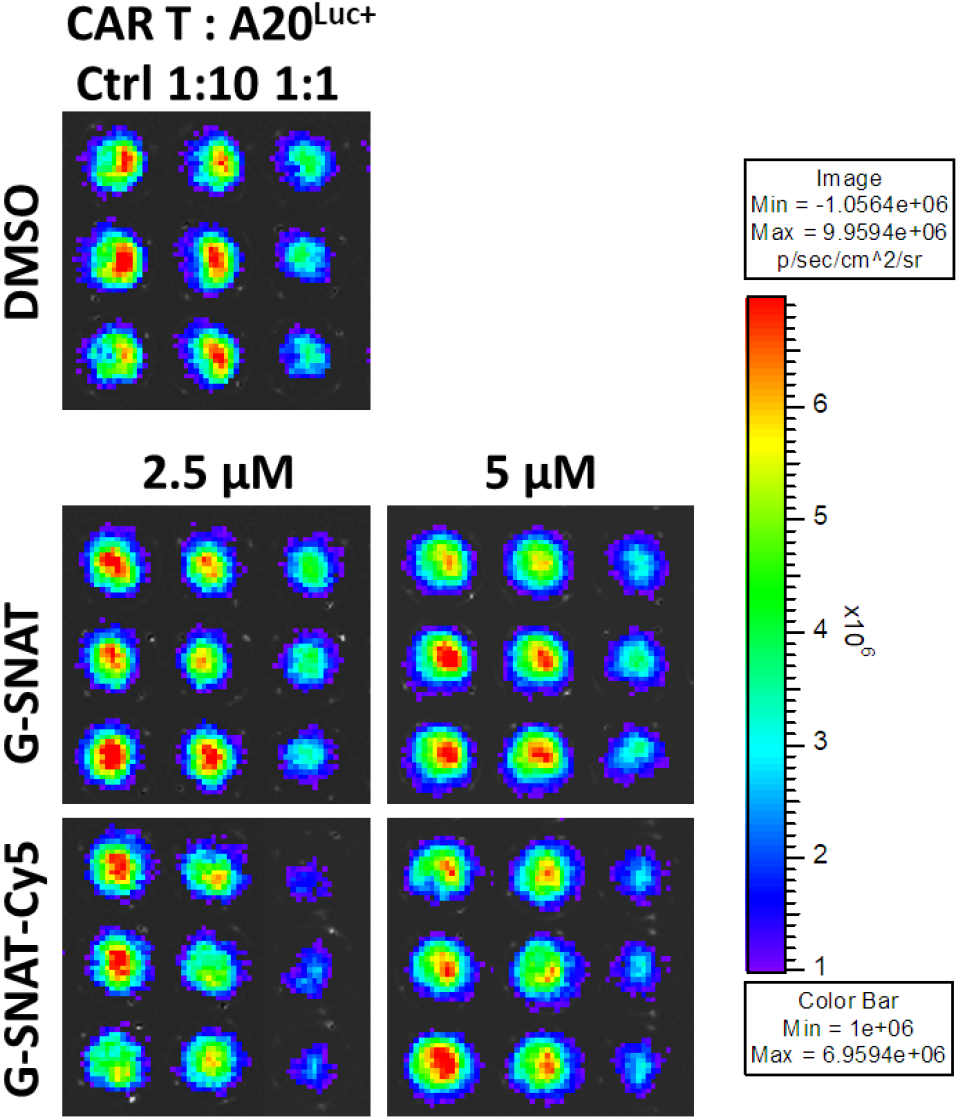
Bioluminescence assay confirmed the cytotoxic function of CD19-28ζ CAR T cells in the presence of G-SNAT or G-SNAT-Cy5. Bioluminescent intensity of target A20^Luc+^ cells cocultured without or with effector CD19-28ζ CAR T cells at 1:10 and 1:1 effector-to-target (E:T) ratios in the presence of G-SNAT, G-SNAT-Cy5 with indicated concentration or solvent (DMSO, 1%) overnight in 96-well plate. D-luciferin (300 μg/mL) was added before scan with an IVIS optical imager.

**Fig. S8.**
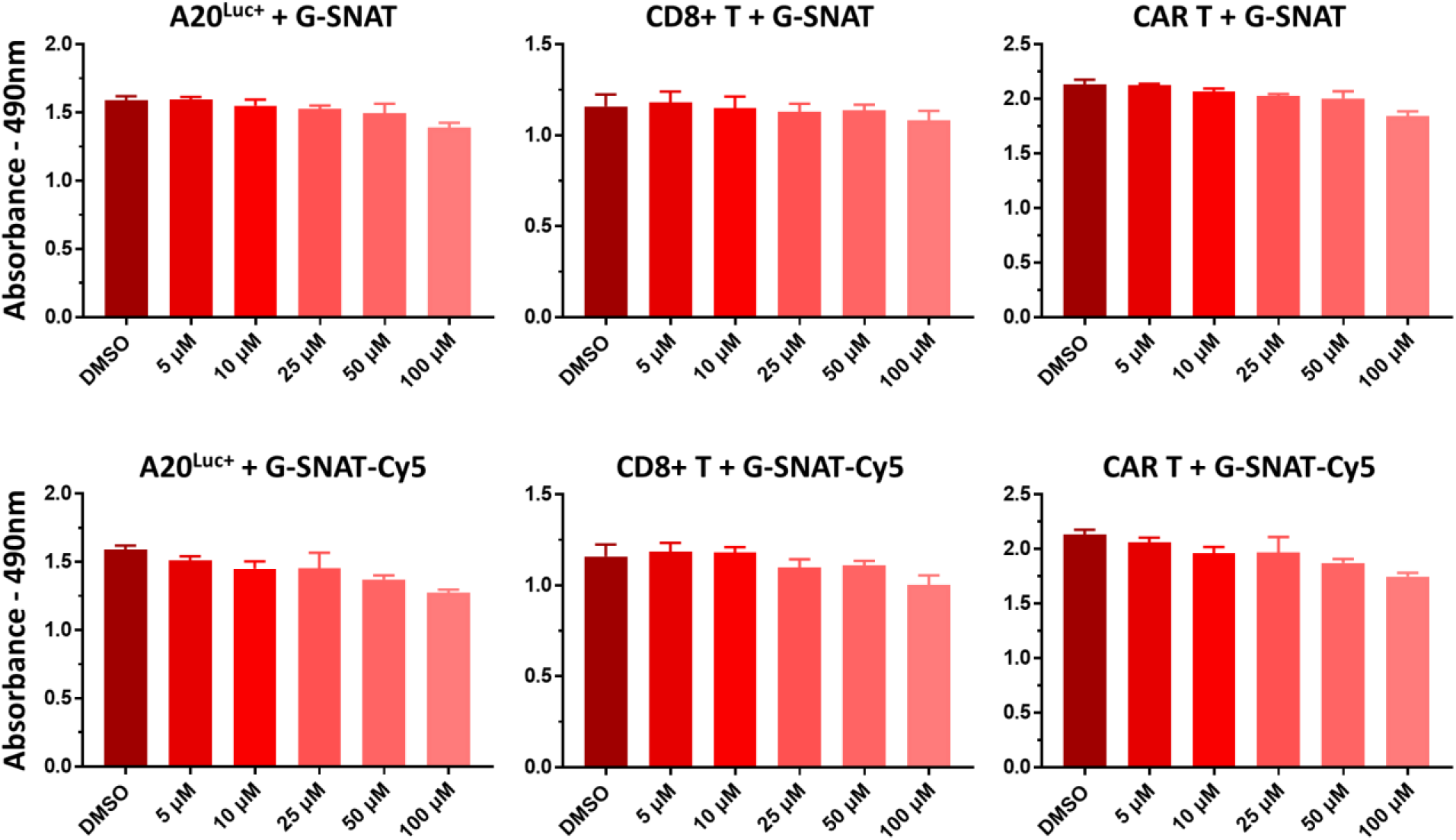
Cell viability study of G-SNAT and G-SNAT-Cy5. Viability assay (MTS) with G-SNAT, G-SNAT-Cy5 and solvent (DMSO, 1%) treated A20^Luc+^, CD8+ T and CAR T cells at 37 °C for 3.5 h. Absorbance at 490 nm was measured with MTS tetrazolium compound.

**Fig. S9.**
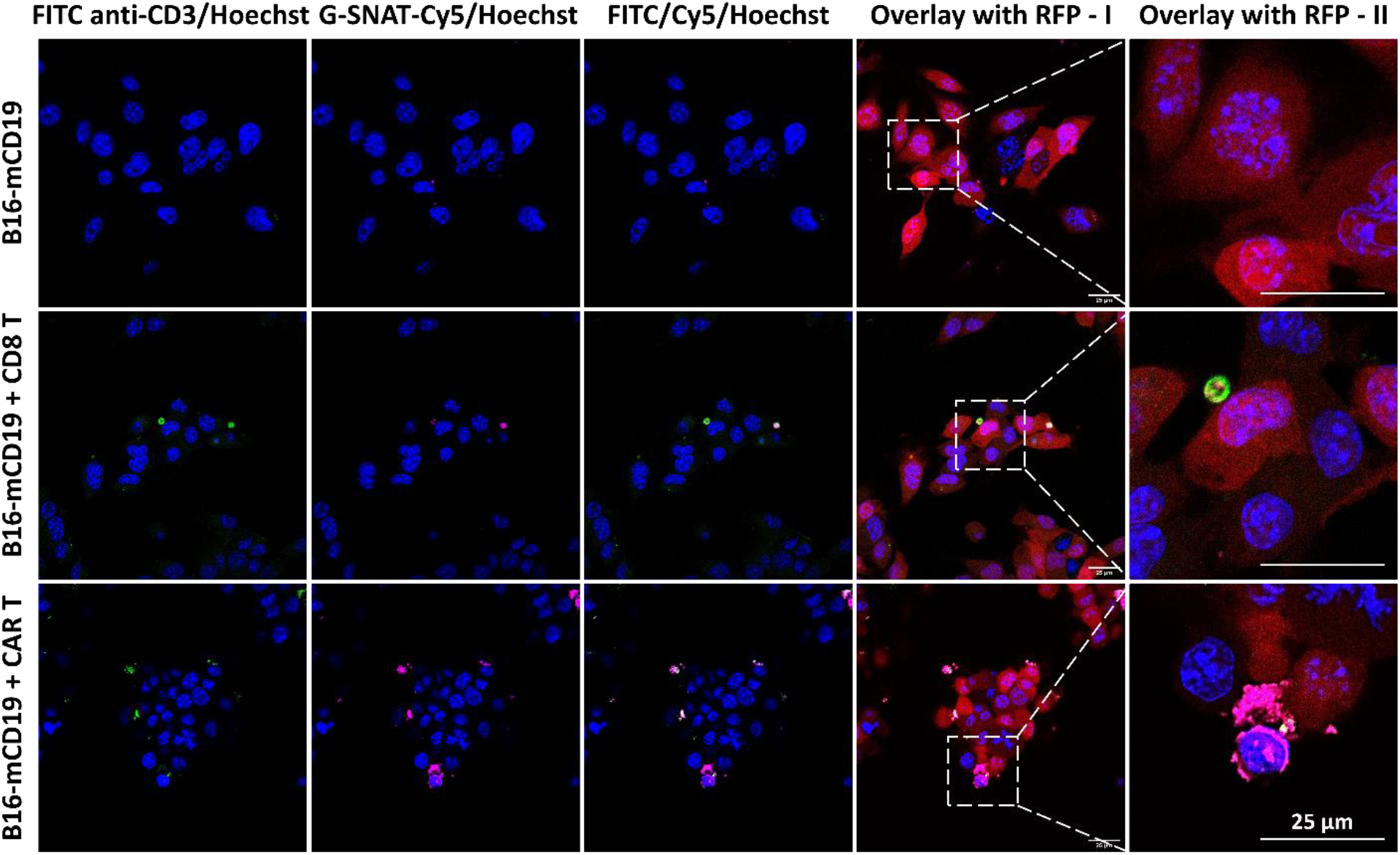
Confocal imaging of B16-mCD19 treated with activated untransduced CD8+ or CD19- 28ζ CAR T cells. Activated untransduced CD8+ or CAR T cells were added to a single layer of B16-mCD19 cells at 2 to 1 ratio in the presence of G-SNAT-Cy5 probe (5 μM) for 3.5 h. Cells were gently washed, stained with Hoechst (blue, Ex390/Em440) and FITC conjugated CD3 antibody (green, Ex488/Em520), fixed and mounted for confocal microscope to show the stacked image. Magenta (Ex650/Em670) represent cyclized and aggregated G-SNAT-Cy5. Red represents RFP (Ex550/Em580). Scale bar indicates 25 μm.

**Fig. S10.**
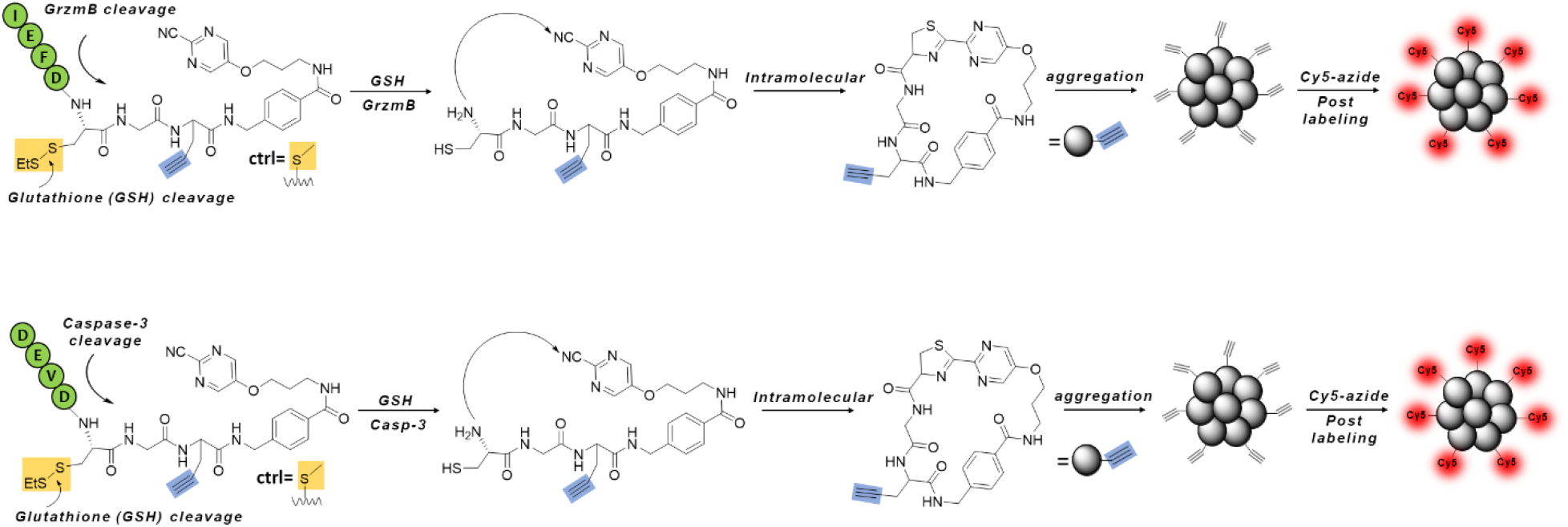
Illustration of the post-click labeling of grzm B activated **G-SNAT** (upper) and caspase- 3 activated **C-SNAT4** (lower) nanoaggregation by Cy5 azide.

**Fig. S11.**
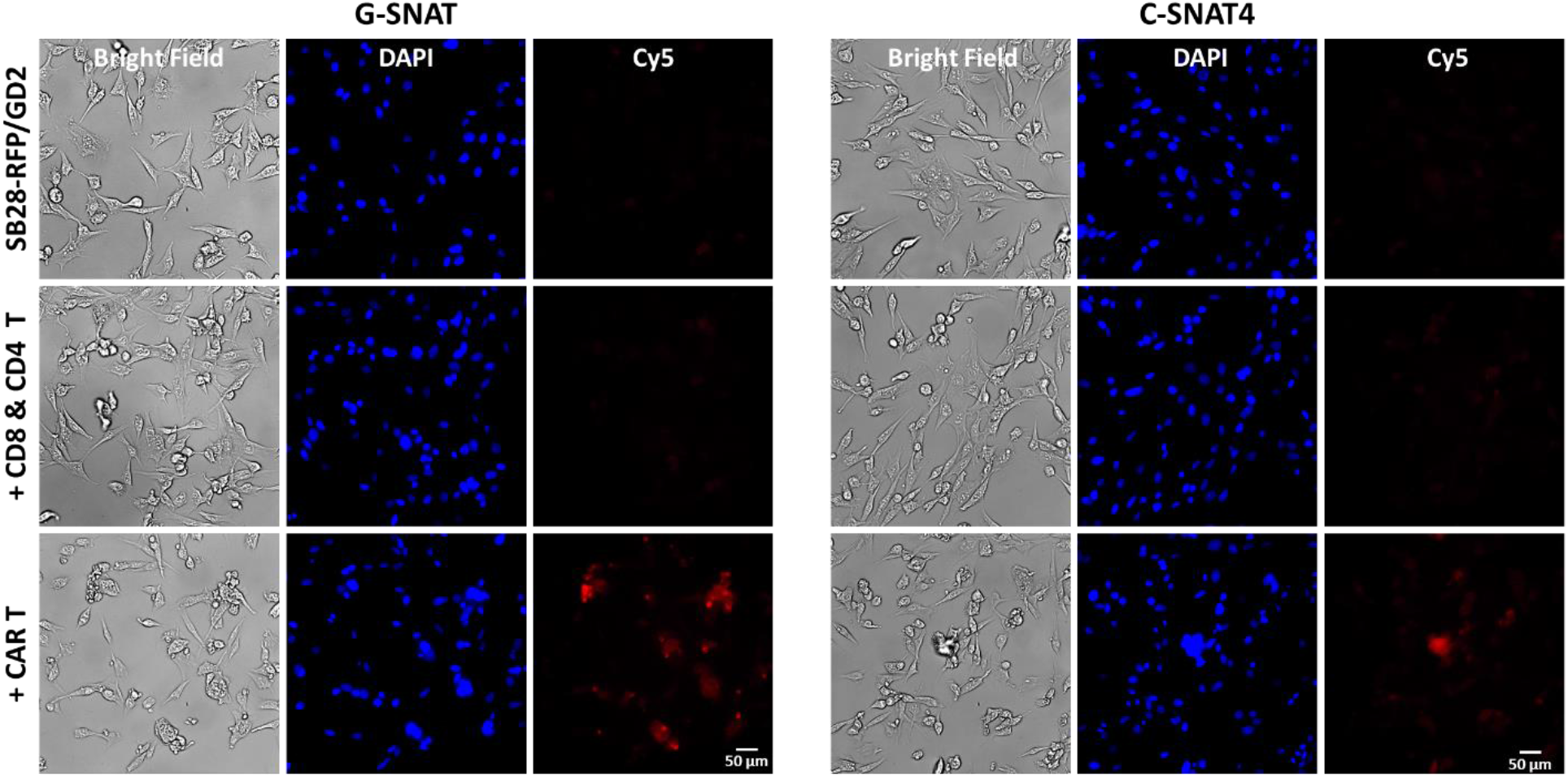
Imaging gzmB and caspase 3 activity in GD2-4-1BBζ CAR T treated SB28-RFP/GD2 cells with G-SNAT-Cy5 and C-SNAT4-Cy5. GD2-4-1BBζ CAR-T cells were added to a single layer of SB28-RFP/GD2 cells at 2 to 1 ratio in the presence of G-SNAT (left, 20 μM) or C-SNAT4 (right, 20 μM) for 2.5 h. Cells were gently washed, permeabilized and processed for Cy5-azide post-click labeling. Cells were also stained with DAPI (blue, Ex405/Em460), fixed, and mounted for epifluorescence microscope imaging. Red (Ex650/Em670) represent cyclized and aggregated G-SNAT-Cy5. Scale bar indicates 50 μm.

**Fig. S12.**
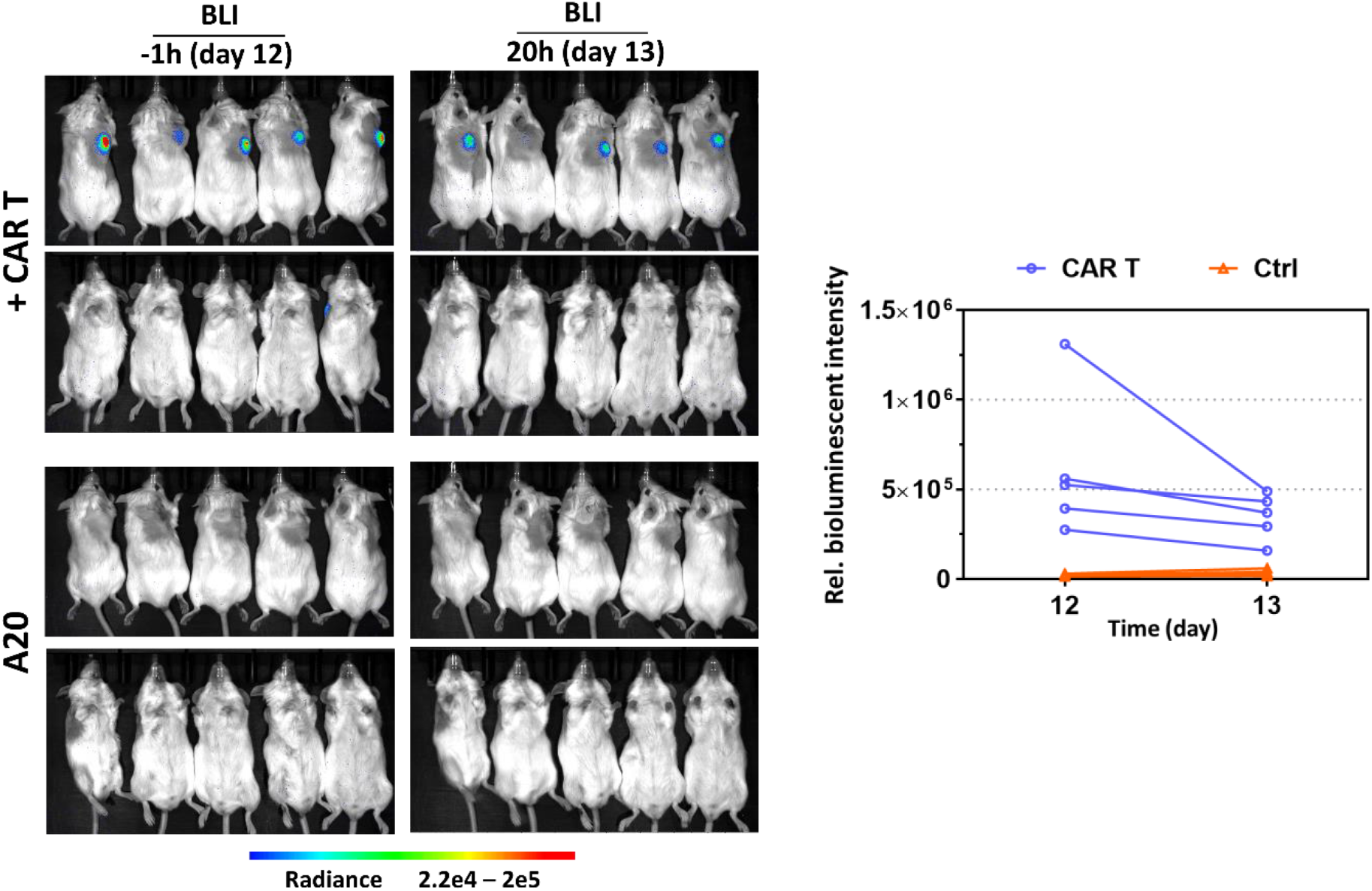
Bioluminescence imaging (left panel) and quantification (right panel) with D-luciferin of A20 implanted (bottom) and CD19-28ζ CAR T^Luc+^ cells (top) treated tumor-bearing mice.

**Fig. S13.**
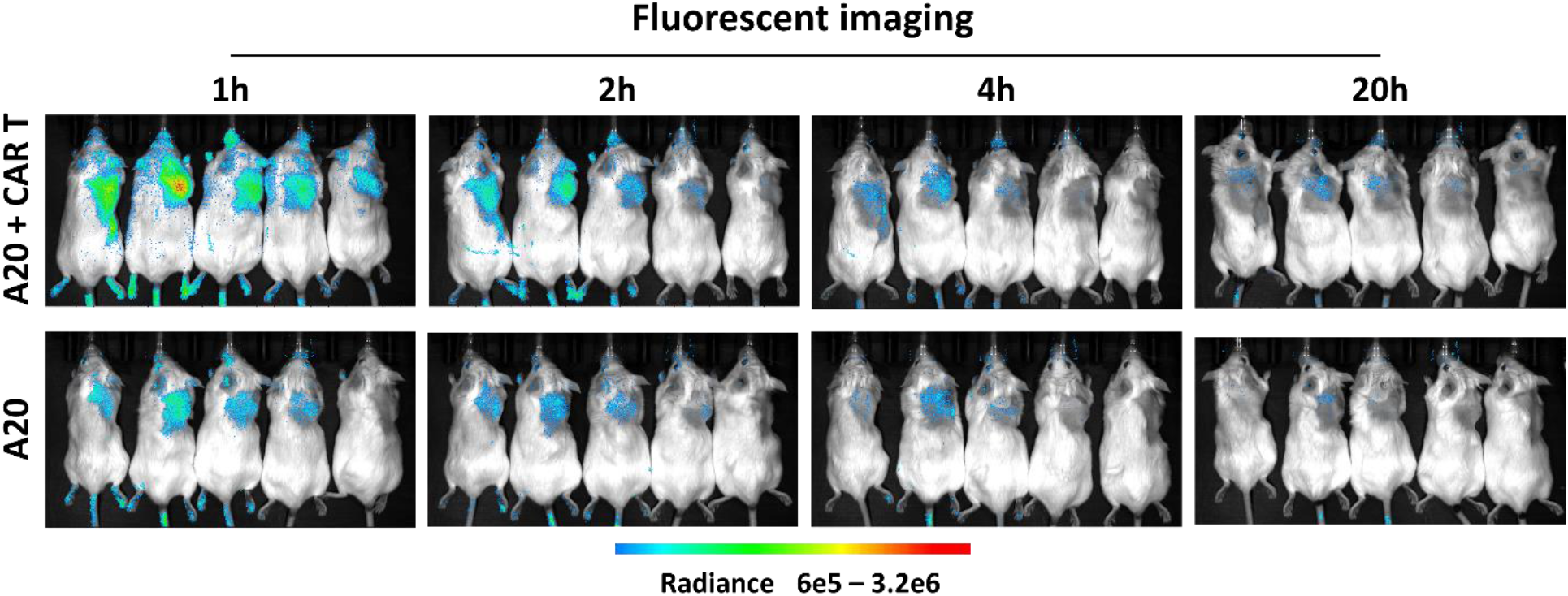
Longitudinal fluorescence imaging with G-SNAT-Cy5 (5 nmol, Ex650/Em670) of A20 implanted (bottom) and CD19-28ζ CAR T^Luc+^ cells (top) treated tumor-bearing mice at day 12.

**Fig. S14.**
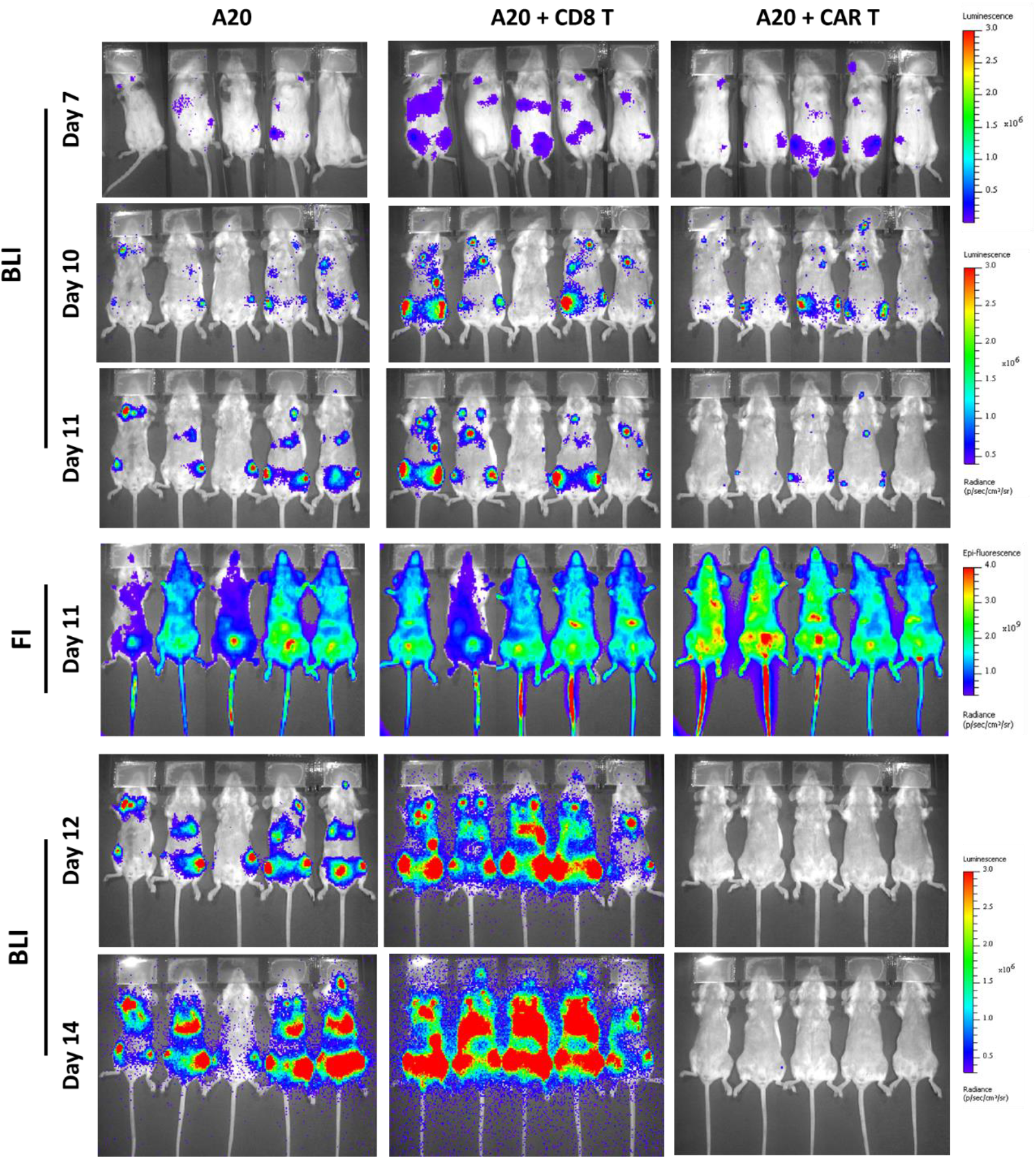
Longitudinal bioluminescence imaging with D-luciferin of A20^Fluc+^ implanted (bottom), activated untransduced CD8+ T cells (middle) or CD19-28ζ CAR T cells (top) treated tumor-bearing mice.

**Fig. S15.**
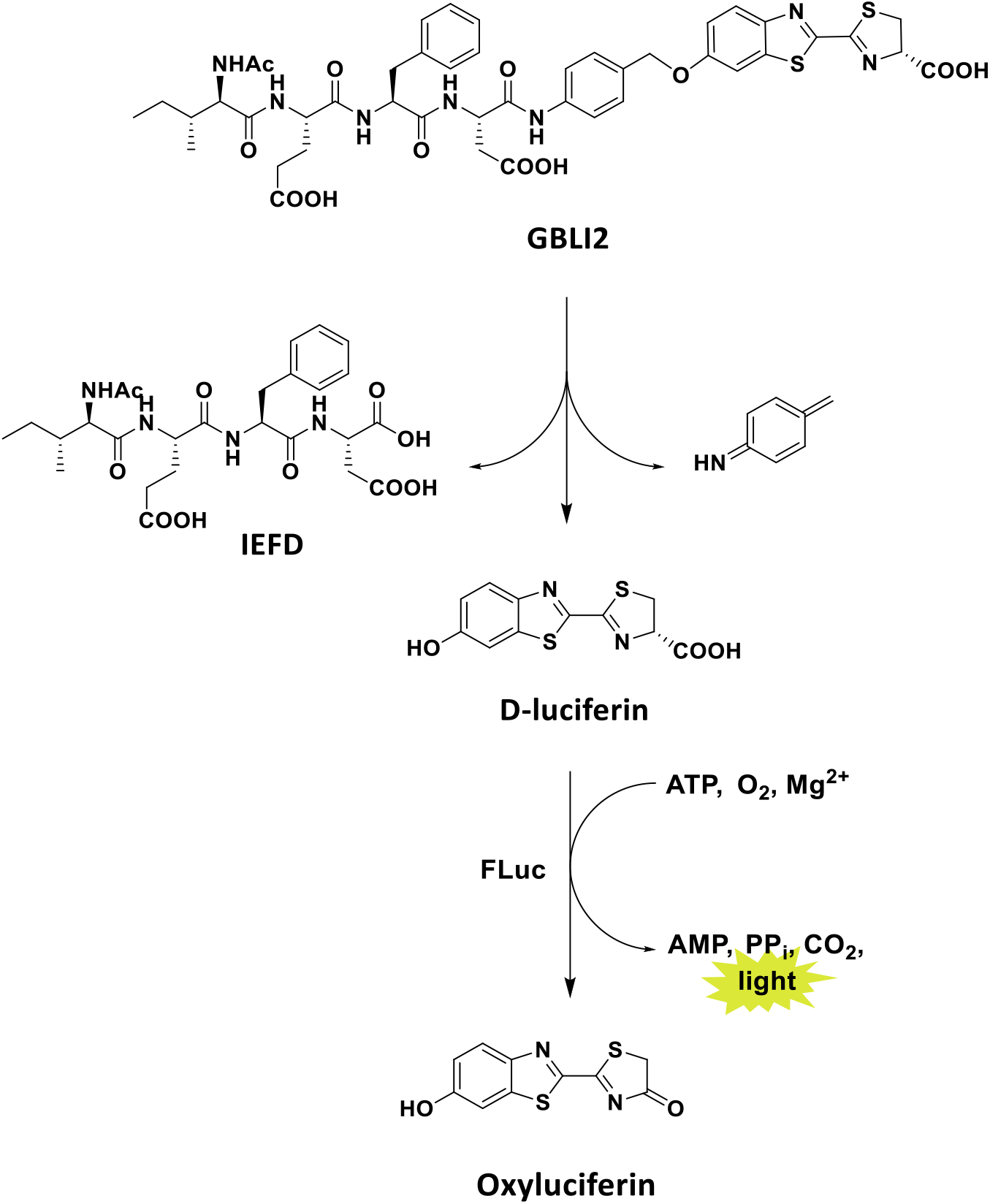
Illustration of the gzmB activated bioluminescent assay with GBLI2. FLuc-firefly luciferase.

**Fig. S16.**
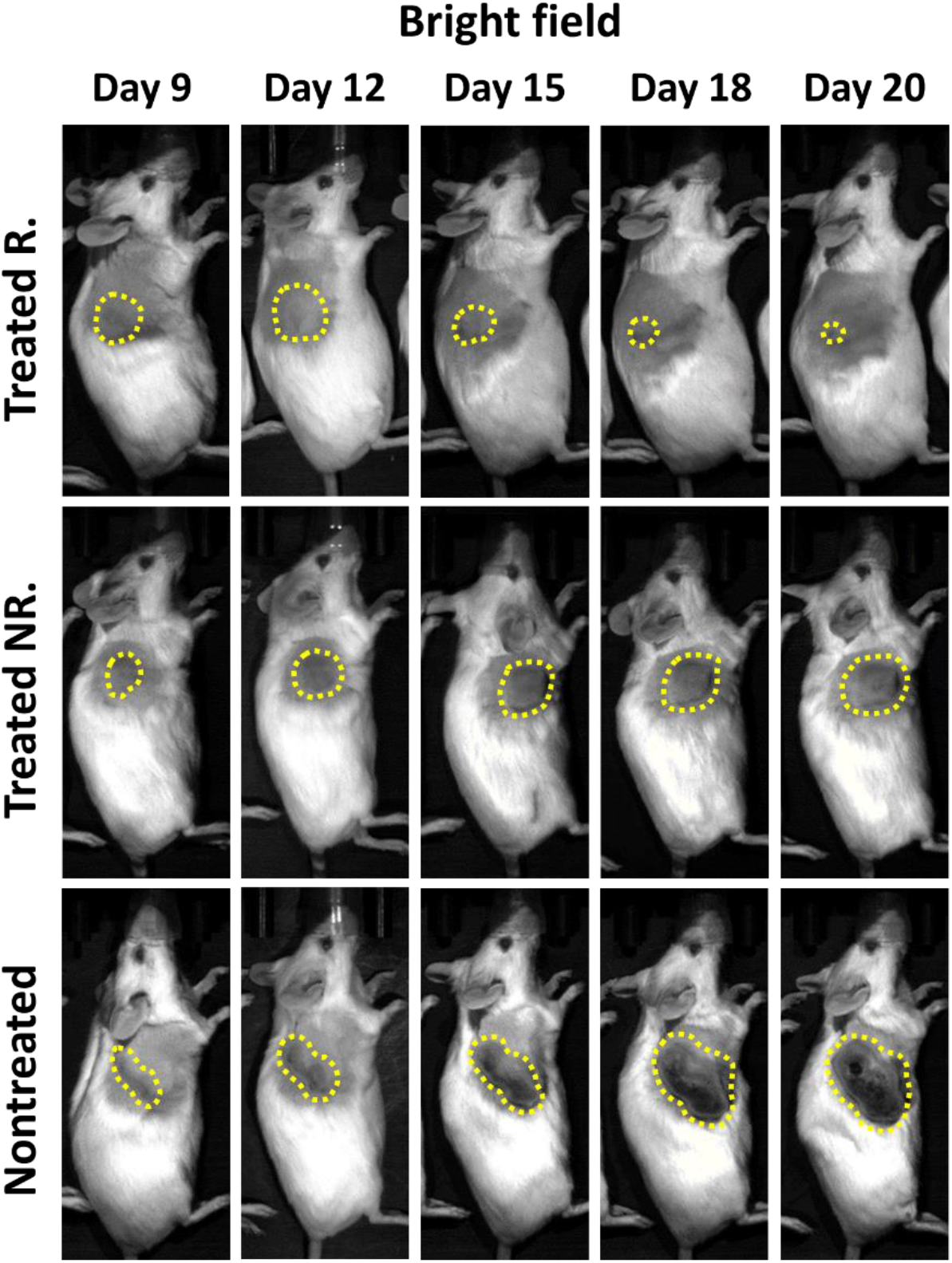
Longitudinal bright field imaging of nontreated (bottom), treated non-responder (middle) and responder (top) groups in Fig. 5D. The yellow circles indicate tumors. Representative mice from each group were shown here.

**Fig. S17.**
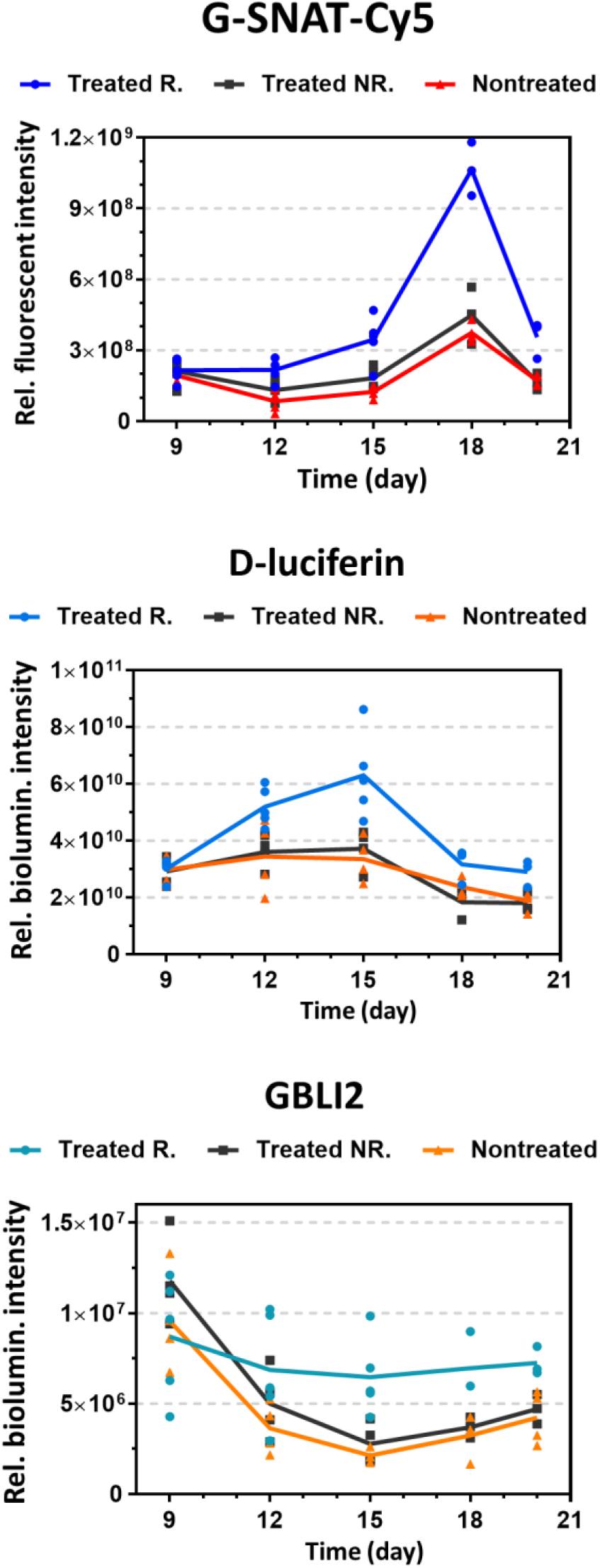
Relative fluorescent or bioluminescent intensity of tumors imaged with G-SNAT-Cy5, D-luciferin or GBLI2 at day 9, 12, 15, 18, and 20 quantified by defining the ROI on tumors.

**Fig. S18.**
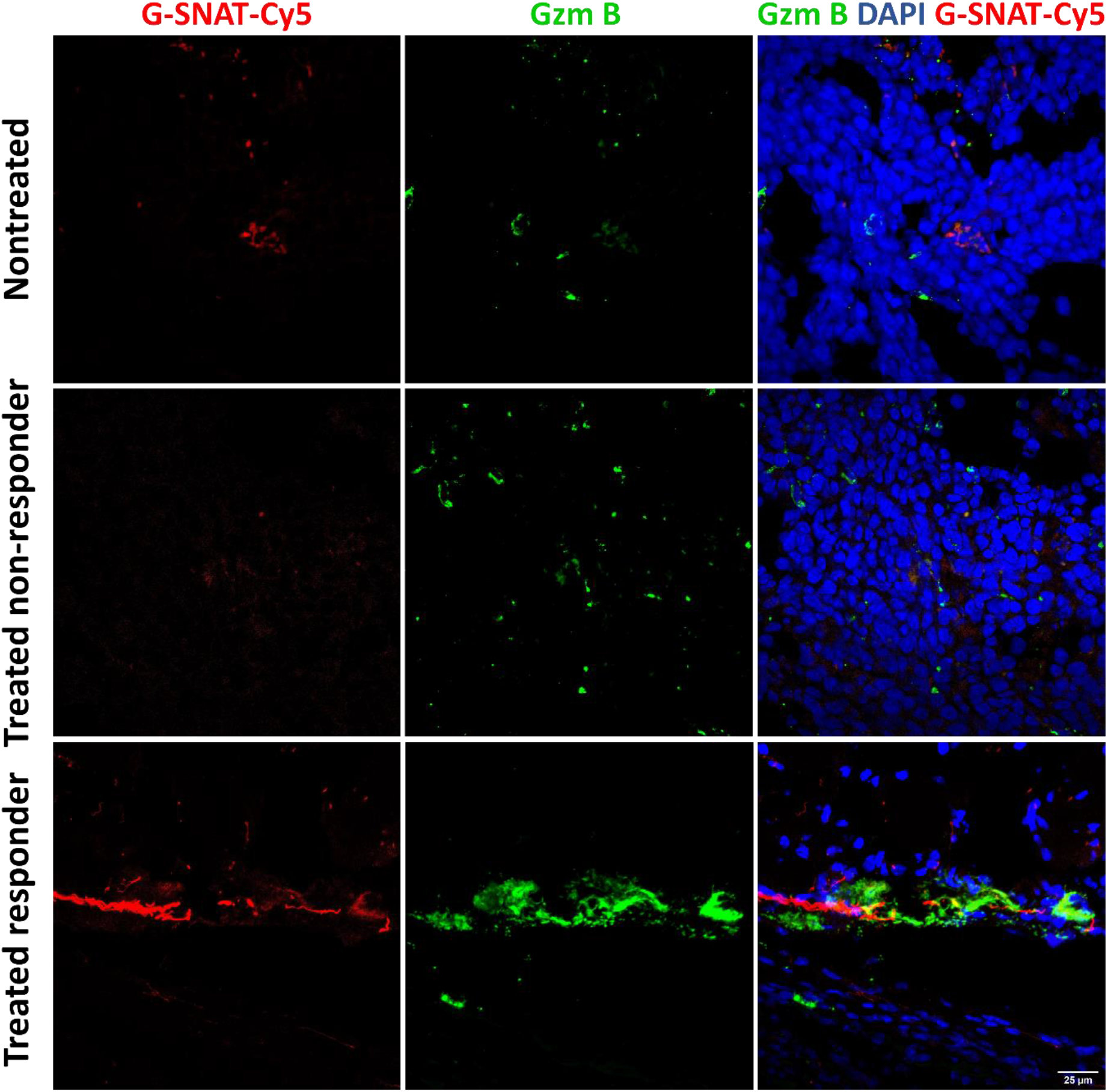
Immunofluorescent staining analysis of the nontreated, treated responded and nonresponded tumors from Fig. 5I. Scale bar indicates 50 μm.

## APPENDIX NMR SPECTRUM

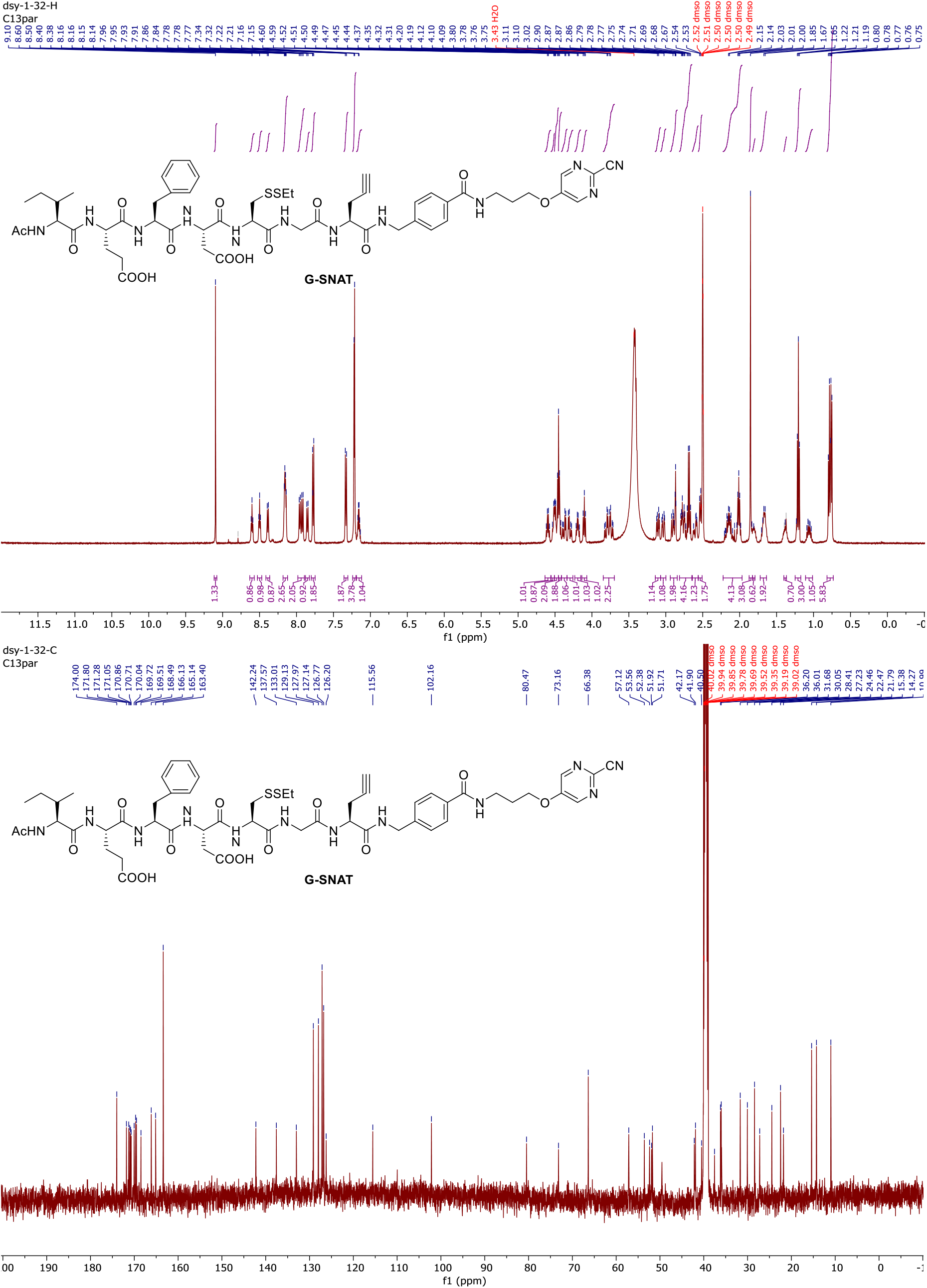

